# A general approach for analysis of physiologically structured population models: the R package ‘PSPManalysis’

**DOI:** 10.1101/2020.06.27.174722

**Authors:** André M. de Roos

## Abstract

1. How environmental conditions affect the life history of individual organisms and how these effects translate into dynamics of population and communities on ecological and evolutionary time scales is a central question in many eco-evolutionary studies.
2. Physiologically structured population models (PSPMs) offer a theoretical approach to address such questions as they are built upon a function-based model of the life history, which explicitly describes how life history depends on individual traits as well as on environmental factors. PSPMs furthermore explicitly account for population feedback on these environmental factors, which translates into density-dependent effects on the life history. PSPMs can thus capture life histories in quite some detail but lead to population-level formulations in terms of partial differential equations that are generally hard to analyse.
3. Here I present a general methodology and a `R` software package for computing how the ecological steady states of PSPMs depend on model parameters and to detect bifurcation points in the computed curves where dynamics change drastically. The package makes specifying the population model unnecessary and only requires a relatively straightforward implementation of the life history functions as input. It furthermore allows for analysing the evolutionary dynamics and evolutionary singular states of the PSPMs based on Adaptive Dynamics theory.
4. Given the central role of the individual life history in many studies, there is substantial scope for using the presented methodology in fields as diverse as ecology, ecotoxicology, conservation biology and evolutionary biology, where it has already been applied to problems like the evolution of cannibalism, niche shifts and metamorphosis.

## Introduction

The life history of individual organisms plays a central role in ecology and evolution, determining the demography of populations and thereby their persistence and existence. Together with the interactions with other species it shapes the dynamics of interacting populations and communities and through mutation and selection it leads to evolutionary change in species traits. Methodologies to assess how life history characteristics translate into consequences at the population level, such as population growth rate, are hence a core part of many ecological and evolutionary studies.

Modelling approaches that describe population and community dynamics explicitly on the basis of individual life histories are referred to as structured-population models (Tuljapurkar & Caswell, 1997). Matrix models (Caswell, 2001) are the most common type of structured-population models. Matrix models describe the population dynamics in discrete time, as do integral projection models (IPMs, Ellner, Childs, & Rees, 2016). Structured-population models describing dynamics in continuous time include stage-(Nisbet & Gurney, 1983) and size-structured population models (Sinko & Streifer, 1969; de Roos, Metz, Evers, & Leipoldt, 1990), which both belong to the more general class of physiologically structured population models (PSPMs, Metz & Diekmann, 1986; de Roos, 1997). Although all structured-population models explicitly include a modeled representation of individual life history, they differ in the way they account for this life history (de Roos, 2020). Matrix models and IPMs are primarily data driven. IPMs, for example, are formulated using (non-)linear relationships that result from fitting observational data on the scaling of somatic growth rate and fecundity with individual body size (Rees, Childs, & Ellner, 2014). In contrast, PSPMs (Metz & Diekmann, 1986; de Roos, 1997) are formulated using a function-based model of the individual life history, which also accounts for the effect of environmental variables, such as food availability and predator density, on this life history (de Roos, 2020). For example, individual foraging, growth and reproduction are in many PSPMs described by a dynamic energy budget model for individual energetics (Kooijman, 2010). In turn, PSPMs account for how these environmental variables are impacted by the population as a whole. PSPMs hence capture with more mechanistic detail how individual-level processes, like energetics, together with the interactions of the individual with its environment shape the life history and how the feedback of the entire population on this environment has a density-dependent impact on that life history. PSPMs are therefore especially suited to analyse how particular mechanisms or aspects of the life history or ecology of an individual would affect the population and community dynamics.

The downside of the increased mechanistic detail of PSPMs is their mathematical tractability (de Roos, 2020). Where linear algebra offers a rich tool set to analyse matrix models and IPMs, simple PSPMs are formulated in terms of the more daunting partial differential equations (Metz & Diekmann, 1986; de Roos, 1997). Fortunately, though, a collection of numerical techniques is now available that allows for analysing the ecological and evolutionary dynamics of even fairly complicated PSPMs (Hin & de Roos, 2019a; ten Brink, de Roos, & Dieckmann, 2019; Chaparro Pedraza & de Roos, 2020). The aim of this paper is to provide an introduction to these techniques and to the `R` package ‘PSPManalysis’ implementing them. This toolbox includes techniques for the demographic and steady-state analysis (Diekmann, Gyllenberg, & Metz, 2003; de Roos, 2008) of PSPMs, which also allow for the analysis of evolutionary dynamics of PSPMs, based on the framework of ‘Adapative Dynamics’ (Dieckmann & Law, 1996; Metz, Geritz, Meszéna, Jacobs, & van Heerwaarden, 1996; Geritz, Kisdi, Meszéna, & Metz, 1998). In addition, the ‘PSPManalysis’ package includes the ‘Escalator Boxcar Train’ (de Roos, Diekmann, & Metz, 1992), a numerical integration technique specifically developed for PSPMs.

To use the ‘PSPManalysis’ package it is not necessary to bother with the population-level representation of the model in terms of partial differential equations or the like. The user can concentrate on the life history and the ecology of the individual organisms. The necessary ingredients of the model specification are conceptually straightforward as they include (*i*) the individual state variables (traits), such as age or body size, that determine the life history; (*ii*) the environmental variables, such as food availability or predation pressure, that impinge on and shape the life history; (*iii*) the rates of development, reproduction and mortality dependent on these individual state and environmental variables; (*iv*) how an individual impacts its environment; and (*v*) the conditions determining that the environment of the individuals is in steady-state. The routines implemented in the ‘PSPManalysis’ package take these individual life history ingredients as input and numerically analyse their population-level consequences, both in ecological and evolutionary time.

## Materials and methods

### The individual life history model

To illustrate the analysis of ecological steady states and the evolutionary analysis of PSPMs with the ‘PSPManalysis’ package I use the life-history model described in Chaparro Pedraza and de Roos (2020) as an example. This model is loosely based on the life history of salmon with individuals starting their life in a safe nursery habitat, in which they are protected from predation but suffer from competition for resources. At some point during their life the individuals switch to a more risky growing habitat, where competition for resources is absent, but individuals are exposed to predation mortality. All equations occurring in the model are presented in Table 1, while Table S1 in the supporting information lists all parameters with their default values.

**Table 1:** Model functions of the individual life history model of Chaparro Pedraza and de Roos (2020).

| Function |  | Description |
| --- | --- | --- |
| *Life history functions* |  |  |
| $\gamma(\ell, X) = \begin{cases} \xi \left( \ell_{inf} \frac{X}{K+X} - \ell \right) \\ \xi (\ell_{inf} - \ell) \end{cases}$ | $\ell < \ell_s$<br>otherwise | Growth rate in length |
| $\beta(\ell) = \begin{cases} B_{max} \ell^2 \\ 0 \end{cases}$ | $\ell > \ell_m$<br>otherwise | Fecundity |
| $\mu(\ell, P) = \begin{cases} \mu_1 \\ \mu_2 + \phi \ell^{-d} P \end{cases}$ | $\ell < \ell_s$<br>otherwise | Mortality |
| *Impacts on the environment* |  |  |
| $\alpha(\ell, X) = \begin{cases} I_{max} \frac{X}{K+X} \ell^2 \\ 0 \end{cases}$ | $\ell < \ell_s$<br>otherwise | Foraging on resource in nursery habitat |
| $\varepsilon(\ell) = \begin{cases} \phi \ell^{3-d} \\ 0 \end{cases}$ | $\ell > \ell_s$<br>otherwise | Contribution to predator numerical response |
| *Functions related to the environment* |  |  |
| $g(X) = \rho (X_{max} - X)$ | | Resource turn-over rate |
| $h(B) = B - \mu_p$ | | Predator per-capita growth rate as a function of the numerical response $B$ |

Individuals are in the model characterised by their length *ℓ* and their population, to which I will refer to as the ‘consumer’ population, is hence length-structured. Migration to the growing habitat and maturation occur on reaching threshold body sizes, at length *ℓ* = *ℓ*_*s*_ and *ℓ* = *ℓ*_*m*_, respectively. Feeding on resources, growth in body size and reproduction are in the model described by the dynamic energy budget (DEB) model developed by Jager, Martin, and Zimmer (2013). In the nursery habitat the consumers compete for a shared resource *X* at a rate that is proportional to their squared length and to the scaled functional response value *X/*(*K* + *X*) (Table 1). In the growth habitat competition is assumed negligible and individuals have *ad-libitum* food. The DEB model predicts individuals under constant resource densities to grow in length following a vonBertalanffy growth curve with ultimate body length equal to *ℓ*_*in f*_ *X/*(*K* + *X*) and *ℓ*_*in f*_ in the nursery and growth habitat, respectively (Table 1). Reproduction only occurs after individuals have migrated to the growth habitat since *ℓ*_*s*_ *< ℓ*_*m*_. Following the DEB model adult fecundity is proportional to squared individual length.

Consumer mortality in the nursery habitat is assumed constant, while in the growth habitat mortality is negatively size-dependent, proportional to *ℓ*^−*d*^ (Table 1). In Chaparro Pedraza and de Roos (2020) size-dependent mortality in the growth habitat is assumed non-dynamic. Here I assume it to be proportional to the density of a dynamic, unstructured predator population that forages on consumers of different length following a linear functional response and that experiences a constant mortality rate *µ*_*p*_. In the model this predator population is represented with its scaled density, which incorporates the (constant) conversion efficiency between ingested biomass of consumers and the predator’s numerical response, indicated with *B* (Table 1). The contribution of individual consumers to predator intake equals the product of their vulnerability to predation and their biomass *ℓ*^3^. Finally, following Chaparro Pedraza and de Roos (2020) turn-over of the resource in nursery habitat in the absence of consumers is described by a semi-chemostat growth equation.

This tritrophic interaction between a resource and a size-structured consumer population in a nursery habitat, which goes through a habitat shift during its life history and subsequently supports a specialist predator population in the growth habitat, is fully determined by 3 life history functions, describing development, reproduction and mortality (*γ*(*ℓ, X*), *β* (*ℓ*), and *µ*(*ℓ, P*) in Table 1, respectively), 2 functions describing the impact of differently sized consumer individuals on their environment through foraging on the resource and contribution to predator food intake (*α*(*ℓ, X*) and *ε*(*ℓ*), respectively), and by 2 functions that determine the dynamics of the shared resource in the nursery habitat and the predator in the growth habitat (*g*(*R*) and *h*(*B*), respectively; Table 1).

### General methodology

In the context of PSPMs Metz and Diekmann (1986) introduced the fundamental distinction between the individual and its environment with accompanying state concepts. The crucial aspect of this distinction is that the individual life history is fully determined by the state of the individual in combination with the state of its environment. Given a constant environment all individuals are therefore independent, which implies that in PSPMs all forms of density dependence operate through the environment. As a further consequence, the individual life history functions are the only necessary ingredients for the computation of population equilibrium states. I will shortly present the general methodology for computation of equilibria in PSPMs here using the tritrophic model introduced above, but it should be stressed that it is straightforward to generalise the methodology to far more complex PSPMs (see Diekmann et al., 2003, for a detailed discussion).

Given a constant, equilibrium resource density 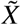 in the nursery habitat the individual length in equi-librium 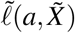 as a function of age *a* and resource density 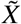 is the solution of the ordinary differential equation (ODE):

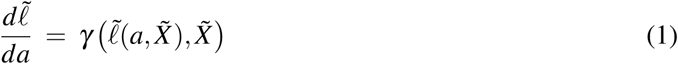

with initial condition 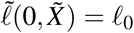. Similarly, denoting the equilibrium predator density in the growth habitat as 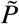, the probability for an individual to survive up to age *a*, which I indicate with 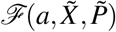 is the solution of the ODE:

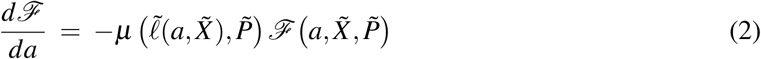

with initial condition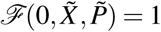. Notice that survival depends on both resource and predator density in equilibrium, as the resource density determines how quickly individual consumers grow and hence how long they experience low mortality in the nursery habitat and the predator density influences their survival in the growth habitat.

The expected reproduction rate by a consumer individual at a particular age equals its fecundity times the probability it survives up to that age. Accumulating these reproductive contributions by integration over all possible ages that individuals can reach results in the following expression for the expected number of offspring produced by a single consumer individual throughout its lifetime, indicated with *R*_0_, as a function of resource density 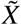 and predator density 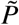:

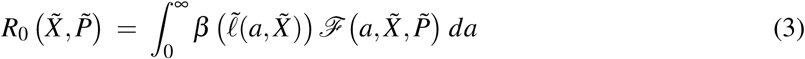

Obviously, the equilibrium state of the size-structured consumer population is determined by the condition 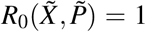, implying that every newborn consumer is expected to just replace itself during life.

For the resource density in the nursery habitat to be in equilibrium the resource turn-over should balance total resource consumption by all consumers in the nursery habitat. The latter equals the product of the amount of resources that an individual consumer is expected to consume during its life time and the consumer population birth rate, which I will indicate with 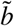. The expected lifetime consumption by an individual is an integral similar to the expression above for *R*_0_ but involving the foraging rate *α*(*ℓ, X*) as opposed to the fecundity *β* (*ℓ*). The steady-state condition for the resource is hence given by the condition:

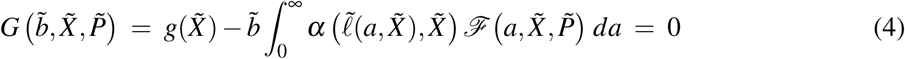

in which the integral represents the expected lifetime resource intake by a single consumer.

Lastly, the predator population in the growth habitat is in steady state when its numerical response *B* equals its per-capita mortality rate *µ*_*p*_. Given the scaling of the predator population density such that its numerical response equals its functional response, the quantity *B* equals the product of the consumer population birth rate 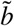 and the expected amount of biomass that a consumer individual during its lifetime contributes to the per-capita food intake rate of the predator. The latter is given by an integral similar to the expression for *R*_0_ in equation (3) but involving the function *ε*(*ℓ*) as opposed to the fecundity *β* (*ℓ*). The steady-state condition for the predator is given by the condition:

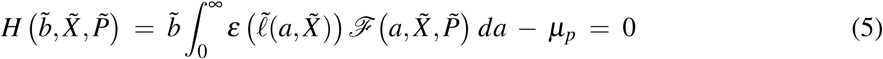

The integral in the above expression represents the expected lifetime contribution by a consumer to the food intake of a single predator.

Even though the ODE (1) for the growth in length is in the current model sufficiently simple to allow for an explicit expression for the length at age 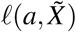 in equilibrium, analytical evaluation of the integrals in the expressions for 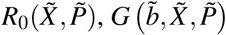 and 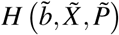 is not possible because of the dependence of consumer mortality on length. Hence, steady states of the PSPM can only be computed by solving the equilibrium conditions (eqs. (3), (4) and (5)) numerically and iteratively for the unknown variables 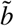, 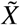 and 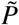. Solving such a system of (non-linear) equations can be achieved by standard methods, such as the Newton-Raphson method (Press, Flannery, Teukolsky, & Vetterling, 1988), but for the fact that it is impossible to derive explicit expressions for the integrals in the functions 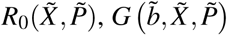 and 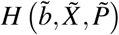 even in a model as simple as the one discussed here. Definitely, the same holds for more complex PSPMs as well.

The key idea to address this issue, originally proposed by Kirkilionis et al. (2001), is to consider the integrals occurring in the equilibrium conditions as a function of the upper limit of the integration and define the following functions:

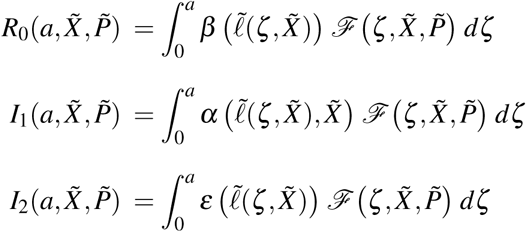

The value of these integrals can then be computed by numerically integrating the system of ODEs:

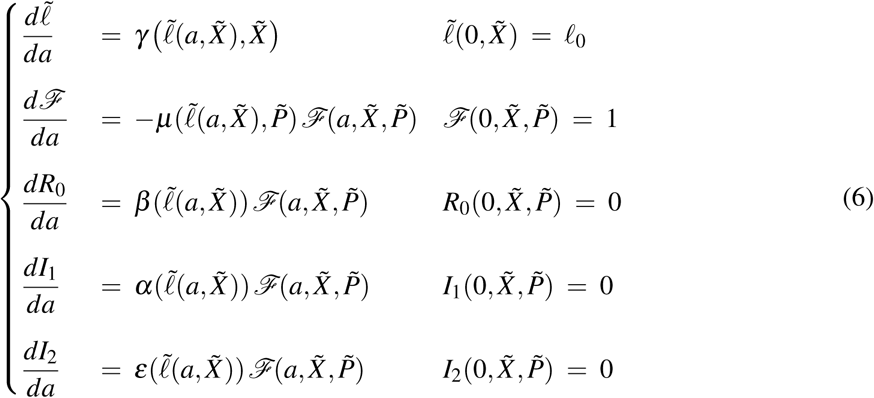

for the length 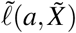 at age *a*, survival 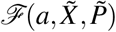, expected cumulative reproduction 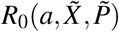, expected cumulative resource ingestion 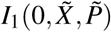 and expected biomass contribution to the predator food intake 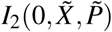 of a consumer individual up to age *a* (The ODEs for these last 3 quantities are derived by differentiating their integral expressions with respect to *a*). Using these quantities the steady-state conditions of the PSPM can be expressed as:

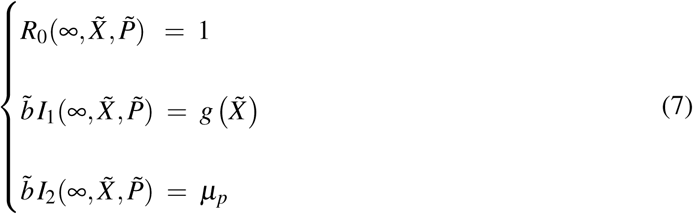

The Newton-Raphson method can be used to solve this system of equations iteratively for the unknown variables 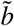, 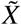 and 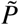, but for every evaluation of these equations the ODEs (6) have to be integrated numerically. This integration in theory has to proceed until infinite age but in practice integration is stopped when the probability to survive 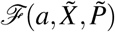 has dropped below some very low value (e.g. 1.0 ·10^−9^).

The methodology discussed above is sufficiently general that it can be applied to a wide range of PSPMs, including those with finitely many individual and environmental state variables and with individuals that are born with finitely many different states at birth (Diekmann et al., 2003). The `R` package ‘PSP-Manalysis’ uses this methodology to compute steady states of PSPMs but also implements pseudoarclength continuation techniques to compute steady state curves as a function of 1 or 2 model parameters (Kuznetsov, 1998, Chapter 10). The results section illustrates how to use this curve continuation approach for model analysis. While computing such curves the ‘PSPManalysis’ package furthermore detects certain bifurcation points, which are points along a curve where the nature of the computed equilibrium undergoes a qualitative change. As illustrated in the results section, such a qualitative change could refer to whether or not a particular equilibrium state can or can not be invaded by a population. For the detection of these bifurcation points the ‘PSPManalysis’ package again uses the techniques and tests presented in Kuznetsov (1998, Chapter 10).

Diekmann et al. (2003) discuss that the approach to compute steady states of PSPMs can also be used to analyse evolutionary dynamics using the theory of Adaptive Dynamics (Dieckmann & Law, 1996; Metz et al., 1996; Geritz et al., 1998). Adaptive dynamics theory explicitly relates evolution by natural selection to population dynamics by considering whether rare mutant phenotypes can invade and take over a resident population. The invasion fitness of such rare mutants is determined by their population growth rate under the environmental conditions imposed by the population with resident phenotype. Because the quantity 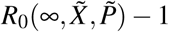 has the same sign as this mutant invasion fitness, it can be used as fitness proxy (Diekmann et al., 2003; Durinx, Metz, & Meszéna, 2007). Therefore, the sign of the selection gradient is determined by the derivative of 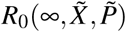 with respect to a model parameter representing a life history trait. I will focus below on the length to switch to the growth habitat *ℓ*_*s*_. Endpoints of evolution in *ℓ*_*s*_, also referred to as *evolutionarily singular strategies* or ESSs then satisfy the condition

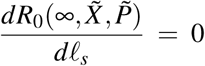

which implies that the invasion fitness reaches a maximum or minimum and the selection gradient vanishes for the given set of environmental conditions 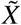 and 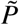. The characteristics of the ESS can be determined on the basis of second-order derivatives of 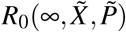 with respect to the life history parameter as explained in detail by Geritz et al. (1998). Furthermore, the fitness gradient 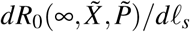 also determines the rate at which the life history trait *ℓ*_*s*_ changes over evolutionary time following the ‘canonical equation of Adaptive Dynamics’ derived by Dieckmann and Law (1996).

Detailed introductions to the theory of adaptive dynamics are found in Dieckmann and Law (1996), Metz et al. (1996), Geritz et al. (1998), Durinx et al. (2007), and Lion (2018). The ‘PSPManalysis’ package implements the techniques and conditions from adaptive dynamics discussed in these publications to locate ESSs and to identify their properties; for example, by classifying them as a ‘continuously stable strategy’, which refers to an ESS to which the life history trait evolves and is furthermore evolutionary stable, or as ‘branching point’, where evolutionary branching or diversification can occur (see Geritz et al., 1998, for details). For this purpose the ‘PSPManalysis’ package computes the first and second-order derivatives of 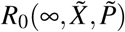 with respect to a life history parameter through numerical differentiation.

### Model implementation for ‘PSPManalysis’

The life history model presented in Table 1 has to be implemented in an `R` script to be analysed with the ‘PSPManalysis’ package. The implementation requires the specification of 3 vectors and 4 functions (see Table 2). The vectors define the model dimensions, the environmental state variables and the model parameters, whereas the functions define the right-hand side of the ODEs (6), their starting values and the boundaries of the different stages in the life history as well as the last two of the conditions (7) that determine the steady states of the environmental variables. In the Supporting Information this implementation of the example model in `R` is discussed in more detail.

**Table 2:**
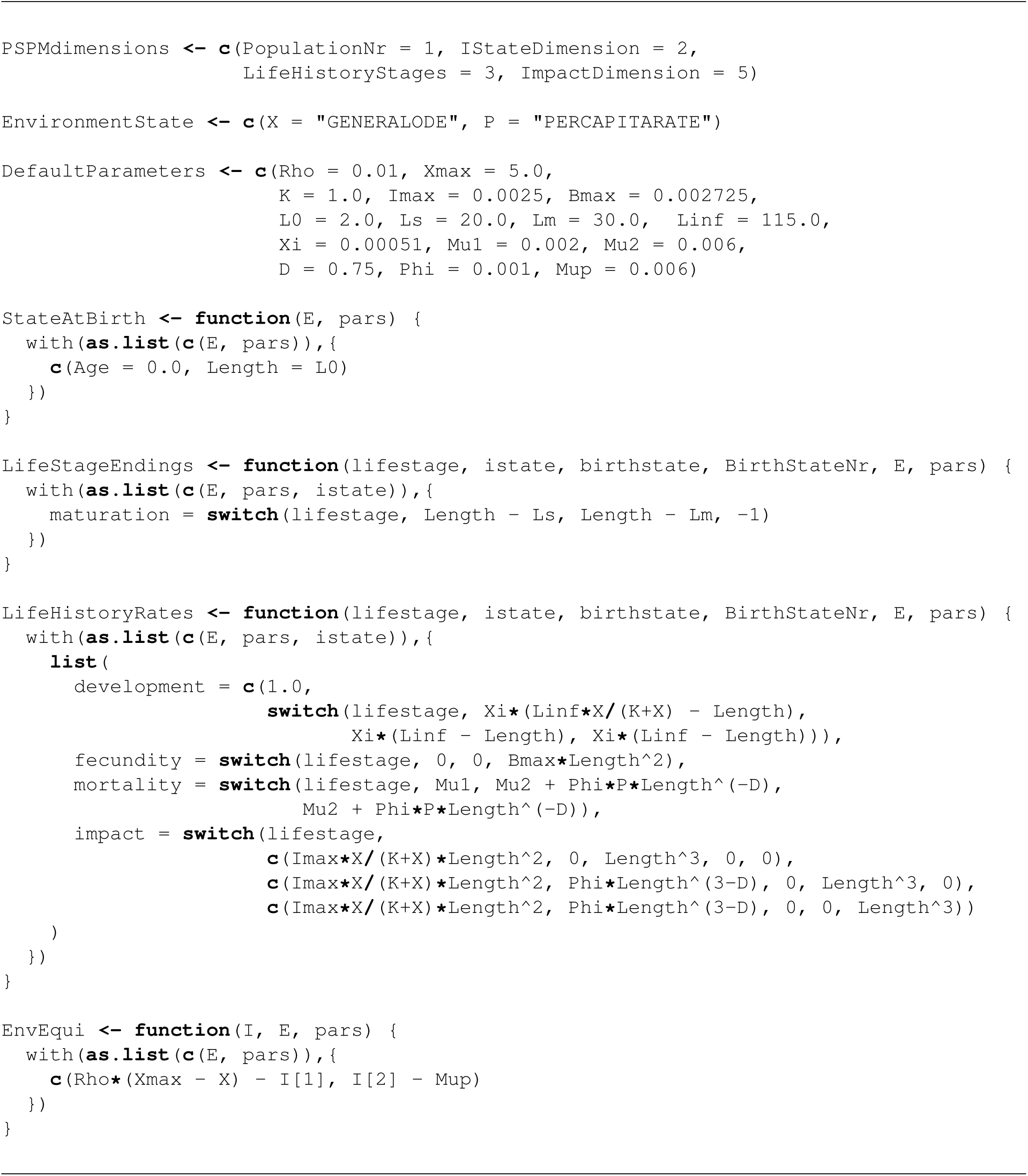
The implementation in `R` of the individual life history model of Chaparro Pedraza and de Roos (2020) for the analysis with the ‘PSPManalysis’ package.

Although the life-history model to be analysed can be specified in `R`, the entire ‘PSPManalysis’ package is written in the `C` language. The package is designed in such a way that implementation of the life history model in `C` is also possible and in fact preferable. The implementation of the example model used in this paper in the `C` language is included in the Supporting Information to this paper as a `C` header file (with extension `.h`). The implementation in `C` has a similar, self-explanatory layout as the model implementation in `R` shown in Table 2 and adaptation of the implementation for another PSPM should only require basic knowledge of the `C` language. Specifying the model definition in `R` may be more straightforward, but comes at a substantial computational cost: Using the model implementation in `C` rather than the `R` implementation decreases the computational time for the results presented in the next section with a factor 40-50.

## Results

### Bifurcation analysis of ecological steady states

The main purpose of the ‘PSPManalysis’ package is to compute the steady states of a PSPM as a function of one of the model parameters, which I from here on refer to as the bifurcation parameter. This not only requires the ‘PSPManalysis’ package implementing the methodology, but also a strategy to execute the computations in a comprehensive manner. Figure 1 shows the equilibrium bifurcation curves of the example model as a function of *X*_*max*_. The figure itself was produced using basic plotting commands in `R` on the basis of data computed with the main function called `PSPMequi` provided by the ‘PSPManalysis’ package. A total of 3 computational steps were needed to generate the data for Figure 1 with the ‘PSPManalysis’ package. Here I discuss these computational steps in broad terms, focussing on the computational strategy rather than on the relevant `R` commands, which are discussed in more detail in the supporting information.

**Figure 1:**
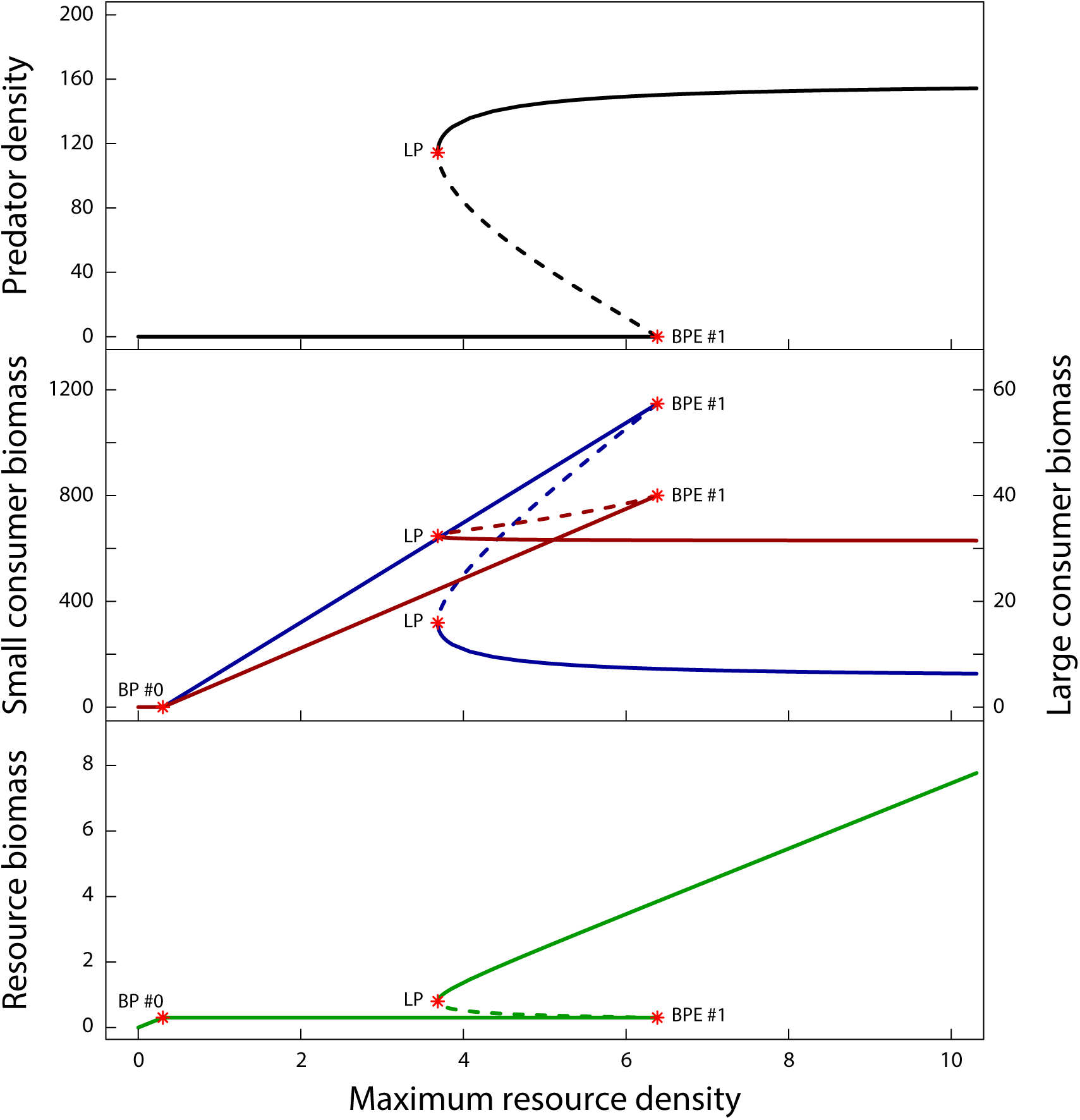
Steady state densities of the unstructured predator population (*top panel*), the length-structured consumer population in the nursery and growth habitat (*middle panel*) and the basic resource (*lower panel*) in the example model (see Table 1) as a function of the maximum resource density *X*_*max*_. All other parameters have default values (Table S1 in the supporting information). See the main text for details about the bifurcation points labeled `“BP #0”` (branching point for structured population with index 0), `“BPE #1”` (branching point for the environmental variable with index 1) and `“LP”` (limit point). Solid lines represent possibly stable equilibria, dashed lines represent saddle points. The curve sections with unstable resource-only and consumer-resource steady states that can be invaded by the structured consumer and unstructured predator, respectively, have been omitted for clarity.

Key to the computation of steady state curves as a function of model parameters is a good starting point. For the example model we can start the computations at the trivial steady state 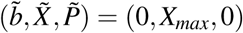, which will be the only steady state for maximum resource densities close to 0, because consumers then do not encounter sufficient resources to reach the length at maturation (note the maximum length equal to *ℓ*_*in f*_ *X/*(*K* + *X*)). As a first step to generating the data for Figure 1 this resource-only equilibrium with zero density of both the length-structured consumer and predator was computed for increasing values of the bifurcation parameter *X*_*max*_, starting from *X*_*max*_ = 0.1. The result is the curve section with increasing equilibrium densities 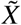 at low values of *X*_*max*_ in Figure 1. This computational step would be superfluous as the value of this steady state is known exactly as 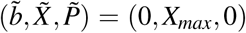, but for the fact that the function `PSPMequi` in the ‘PSPManalysis’ package with which this computational step is carried out, locates a bifurcation point along the curve. It labels this bifurcation point with the string `“BP #0”`, indicating that this bifurcation point represents a branching point (also called transcritical bifurcation point (Kuznetsov, 1998)) for the structured population with index 0 (the consumer population; because the ‘PSPManalysis’ package is written in `C`, indices conform to the `C` convention, in which the first element of a vector has index 0 rather than index 1 such as in `R`). From the output of the function `PSPMequi` it can be inferred that for *X*_*max*_ values to the left of this branching point consumers can not establish themselves in the computed, resource-only equilibrium, because their lifetime reproductive output *R*_0_ is below 1, whereas they can for values to the right of it. These steady states to the left and right of the bifurcation point are therefore stable and unstable (saddle points), respectively.

Step 2 of the computations uses the bifurcation point located in the first computational step as starting point to compute the curve of consumer-resource steady states as a function of *X*_*max*_. The resulting curve corresponds to the part of the equilibrium curves shown in Figure 1 with constant resource density, linearly increasing densities of consumer biomass in the nursery and growth habitat and zero density for the unstructured predator. In this curve the `PSPMequi` function detects a branching point (or transcritical bifurcation point) for the environment variable with index 1 (the unstructured predator population), which it labels as `“BPE #1”` (see Figure 1). The consumer-resource steady states to the right of this branching point can be invaded by the unstructured predator population, as indicated by the positive per-capita growth rate that the function `PSPMequi` produces as output. Predators can hence increase in abundance from low densities for higher *X*_*max*_-values. In contrast, predators have a negative per-capita growth rate and can thus not invade the consumer-resource steady state for *X*_*max*_-values to the left of the branching point labeled `“BPE #1”` in Figure 1, which may therefore represent a stable equilibrium. Whether or not these steady states are indeed stable or, alternatively, cycles in consumer and resource abundance occur can only be investigated using numerical studies of the dynamics, because appropriate test functions that detect transitions from stable equilibrium states to limit cycles (occurring at so-called Hopf bifurcation points (Kuznetsov, 1998)) are currently not available for PSPMs.

The last step of the analysis uses the detected branching point for the unstructured predator population to start a computation of the steady states with positive predator density as a function of *X*_*max*_. The result of this computation is a folded curve of steady state values which extends to a minimum just below *X*_*max*_ = 4. This point at which the equilibrium curve doubles back on itself is another bifurcation point called limit point (or saddle-node bifurcation point Kuznetsov, 1998) and labeled by the function `PSPMequi` with `“LP”` (see Figure 1). Ecologically, this minimum value of *X*_*max*_ represents the persistence boundary of the unstructured predator population whereas the branching point detected in step 2 and labeled `“BPE #1”` represents the predator’s invasion boundary. In between the persistence and invasion boundary two steady states are possible: a consumer-resource steady state that can not be invaded by the predator and a predator-consumer-resource equilibrium. On the basis of the general bifurcation theory presented in Kuznetsov (1998) it can be inferred that the part of the predator-consumer-resource equilibrium curve between the two bifurcation points represents unstable equilibrium states (saddle points).

Because they represent the invasion and persistence boundary of the predator, it may be ecologically relevant to assess how the location of the bifurcation points labeled `“BPE #1”` and `“LP”` depends on other model parameters. The `PSPMequi` function therefore also allows for the computation of these bifurcation points as a function of a second model parameter. Figure 2 shows the location of these two bifurcation points as a function of the maximum resource density *X*_*max*_ and the predator mortality rate *µ*_*p*_. For values in the region of parameter space between the two lines in Figure 2 two steady states occur that are potentially stable, a tritrophic steady steady state with predators and a consumer-resource steady state that predators can not invade. This region of bistability is larger at higher rates of predator mortality.

**Figure 2:**
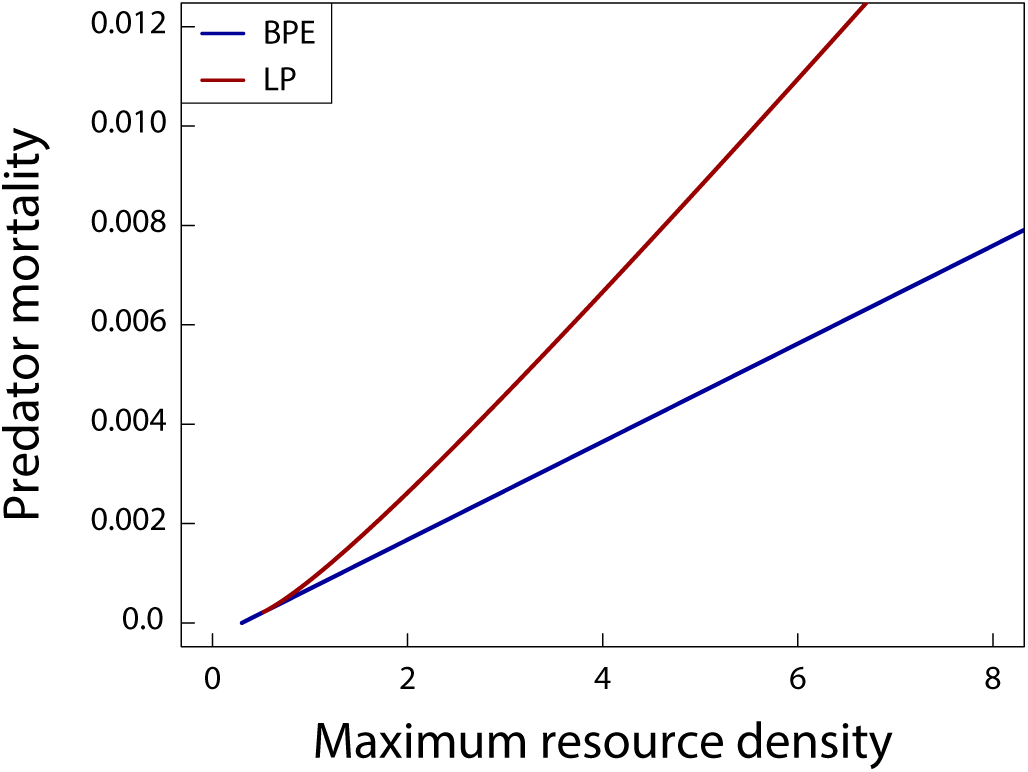
Location of the bifurcation points shown in Figure 1 as a function of the maximum resource density *X*_*max*_ and the predator mortality rate *µ*_*p*_. Only the location of the bifurcation points labeled `“BPE #1”` (branching point for the environmental variable with index 1) and `“LP”` (limit point) are shown, as the location of the bifurcation point labeled `“BP #0”` (branching point for structured population with index 0) is independent of the predator mortality rate *µ*_*p*_. All other parameters have their default values (Table S1 in the supporting information).

### Evolutionary analysis

A particular strength of the ‘PSPManalysis’ package is that it can also be used to calculate evolutionary singular strategies (ESSs) using the framework of Adaptive Dynamics (Dieckmann & Law, 1996; Metz et al., 1996; Geritz et al., 1998). The package therefore allows studying questions about the evolution of life history traits in a context with population and community feedback on the environment in which the evolutionary process takes place. Figure 3 provides as an example the equilibrium bifurcation curves of the example model as a function of the individual length at the habitat switch *ℓ*_*s*_. Because it influences the extent of resource competition and predator vulnerability an individual experiences throughout life, this trait can be expected to be under strong selection. The data shown in Figure 3 have been computed as before with the function `PSPMequi`, which can also produce as output the value of the selection gradient, that is, the derivative 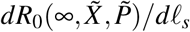 of the lifetime reproductive output *R*_0_ with respect to the life-history parameter *ℓ*_*s*_ (see bottom panel of Figure 3). In the absence of the predator selection for a smaller size at habitat shift occurs, but in the presence of the predator the function `PSPMequi` detects an evolutionary singular state (ESS), which it labels as `“CSS #0”` on the basis of the second-order derivatives of *R*_0_ with respect to *ℓ*_*s*_. The label indicates that the ESS is convergent stable, such that the value of *ℓ*_*s*_ will evolve toward the *ℓ*_*s*_ value of this CSS, while after fixation mutants with slightly different values of *ℓ*_*s*_ will not be able to invade (see Geritz et al., 1998, for further details about the ESS classification).

**Figure 3:**
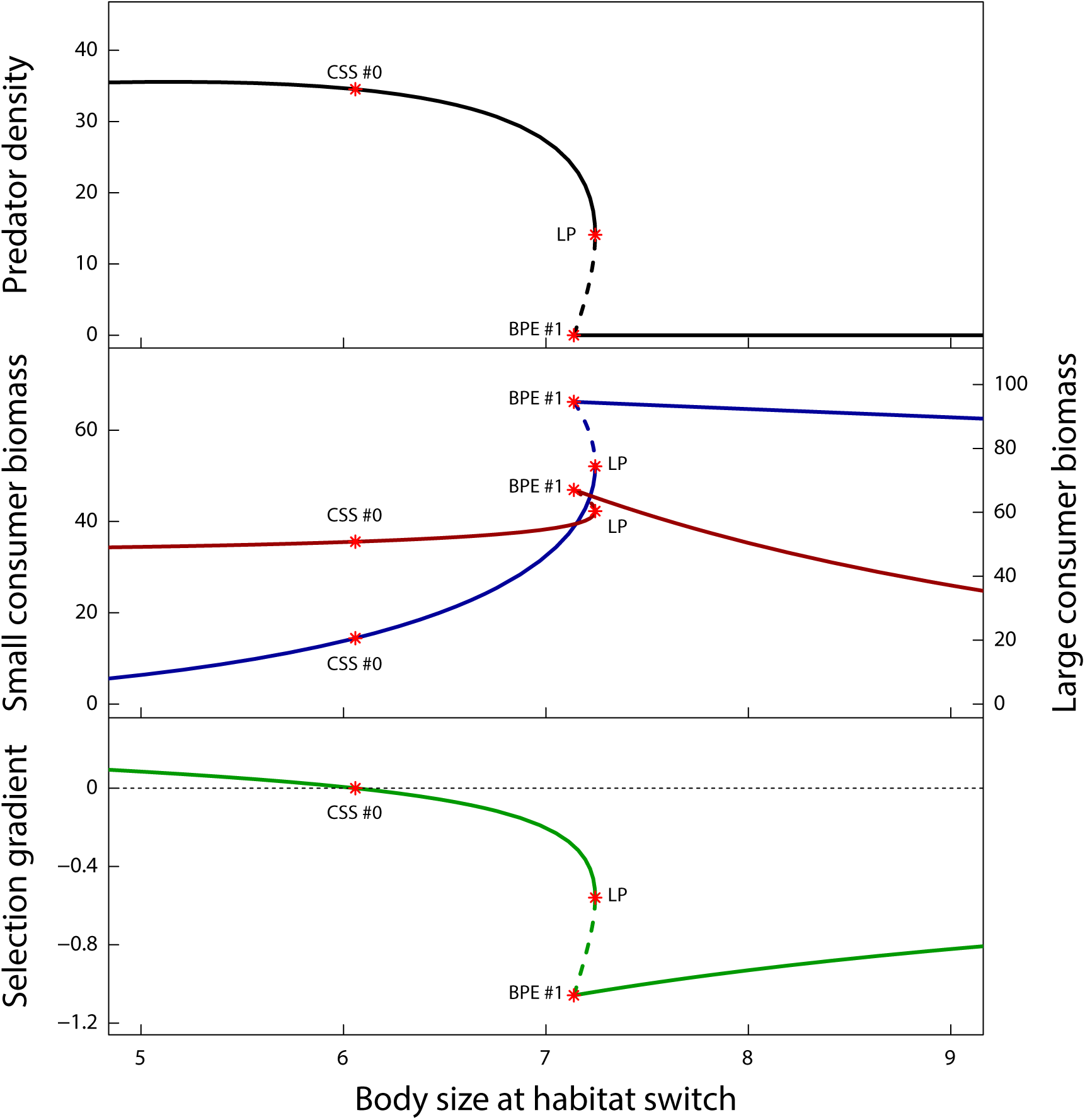
Steady state densities of the unstructured predator population (*top panel*) and the length-structured consumer population in the nursery and growth habitat (*middle panel*), as well as the selection gradient on the individual length at habitat switch *ℓ*_*s*_ in the example model (see Table 1) as a function of this length at habitat switch *ℓ*_*s*_. Maximum resource density *X*_*max*_ = 0.5, all other parameters have their default values (Table S1 in the supporting information). See the main text for details about the bifurcation points labeled `“BPE #1”` (branching point for the environmental variable with index 1) and `“LP”` (limit point) and the evolutionary steady state labeled `“CSS #0”` (convergent stable evolutionary state for structured population with index 0). Solid lines represent possibly stable equilibria, dashed lines represent saddle points. The curve sections with unstable resource-only and consumer-resource steady states that can be invaded by the structured consumer and unstructured predator, respectively, have been omitted for clarity.

Once an evolutionary singular strategy has been detected the function `PSPMequi` can be used to construct a pairwise invasibility plot (PIP; Van Tienderen & de Jong, 1986; Geritz et al., 1998), that is, a graph of the sign of a mutant’s invasion fitness as a function of the life-history trait value of both the resident and the mutant (Figure 4, left panel). Starting from the CSS detected in Figure 3 the function `PSPMequi` was used to compute a boundary with zero mutant fitness as a function of the resident and mutant life-history trait value. This boundary corresponds to the curve separating regions with positive and negative mutant fitness in the PIP shown in Figure 4 (left panel).

**Figure 4:**
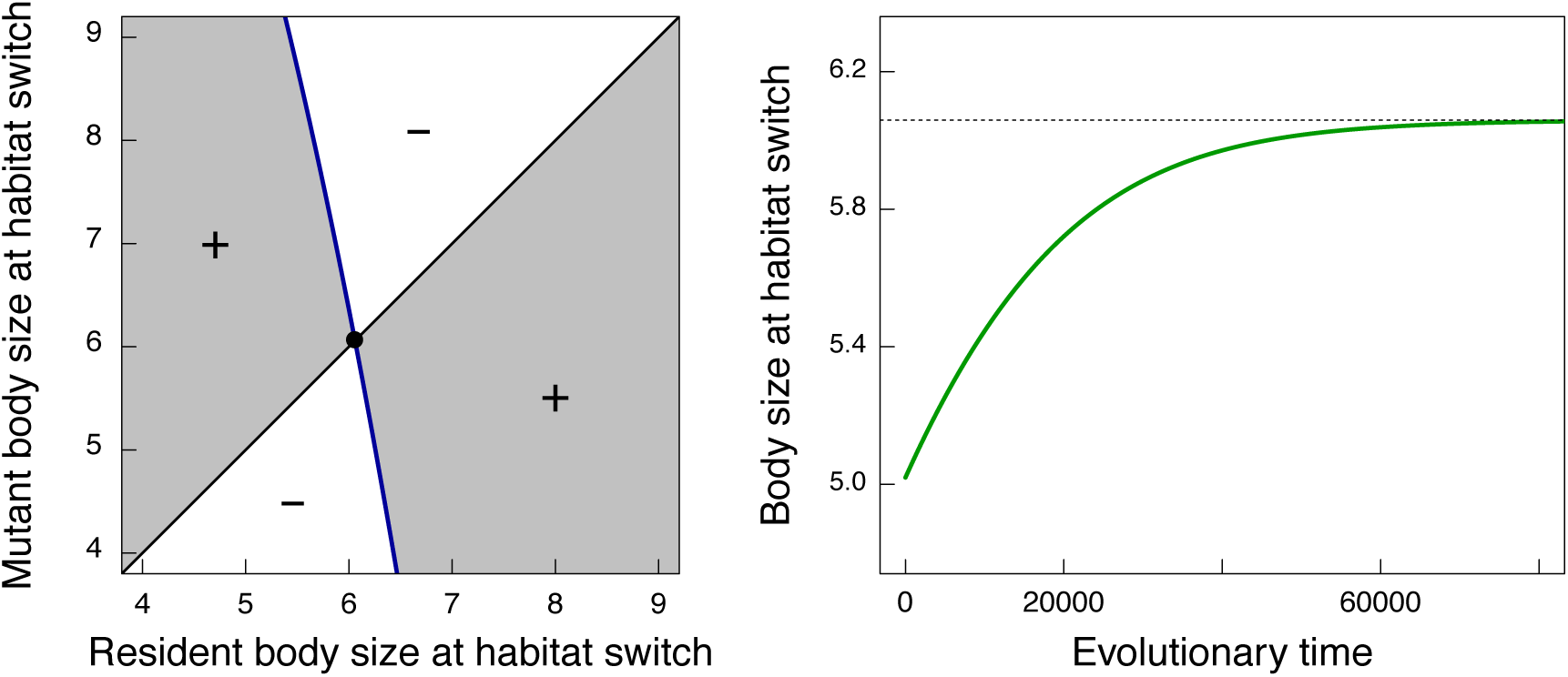
*Left:* Pairwise invasibility plot showing combinations of the resident and mutant value of the length at habitat switch *ℓ*_*s*_ with positive and negative invasion fitness of the mutant. *Right:* Simulation of the dynamics of the evolving value of the length at habitat switch *ℓ*_*s*_ over evolutionary time as predicted by the canonical equation of Adaptive Dynamics (Dieckmann & Law, 1996).

Finally, the ‘PSPManalysis’ package also includes a function `PSPMevodyn`, which can be used to simulate the dynamics of evolving life-history parameters over evolutionary time (Figure 4, right panel). These evolutionary dynamics are described by the canonical equation of Adaptive Dynamics (Dieckmann & Law, 1996). The trajectory of *ℓ*_*s*_ over evolutionary time shown in Figure 4 (right panel) confirms that the individual length at the habitat switch evolves to the value of the convergent stable ESS shown in Figure 3, but since the function `PSPMevodyn` can not simulate combined mutant and resident dynamics, it is not possible to verify whether or not evolutionary branching is possible at this ESS.

The results shown in Figure 3 and 4 were on purpose computed with a low value of *X*_*max*_ = 0.5, as the default value of *X*_*max*_ = 5.0 results in a more complicated, and hence also more intriguing evolutionary outcome of the selection process in *ℓ*_*s*_ (Figure 5). For this value of *X*_*max*_ the range of *ℓ*_*s*_ over which both a consumer-resource equilibrium and a predator-consumer-resource equilibrium occur is more extensive and the evolutionary singular state (classified by the `PSPMequi` function as an evolutionary repellor and labeled `“ERP #0”`) now occurs on the part of the curve representing saddle-node steady states of the predator, consumer and resource. This state is hence ecologically unstable and thus unreachable. As before smaller values of *ℓ*_*s*_ are selected for in the absence of the predator, as this reduces competition for resources among consumers. In the presence of the predator, however, larger values of *ℓ*_*s*_ confer a selective advantage. As a consequence, with predators present evolution will result in larger values of *ℓ*_*s*_ until the limit point is reached around *ℓ*_*s*_ = 22.5, at which point the predator goes extinct and the community collapses to a consumer-resource equilibrium. After predator extinction, the direction of evolution reverses and smaller values of *ℓ*_*s*_ are selected for until the value of this life-history parameter reaches the bifurcation point labeled `“BPE #1”` in Figure 5, below which the predator can once again invade the consumer-resource equilibrium. From Figure 5 it can hence be inferred that over evolutionary time cycling will occur between a consumer-resource steady state and a steady state with predator, driven by the selection process in *ℓ*_*s*_. Simulating the evolutionary dynamics with the `PSPMevodyn` function following the canonical equation of Adaptive Dynamics would, however, not reveal such evolutionary cycling as this function can not switch between the two ecological steady states.

**Figure 5:**
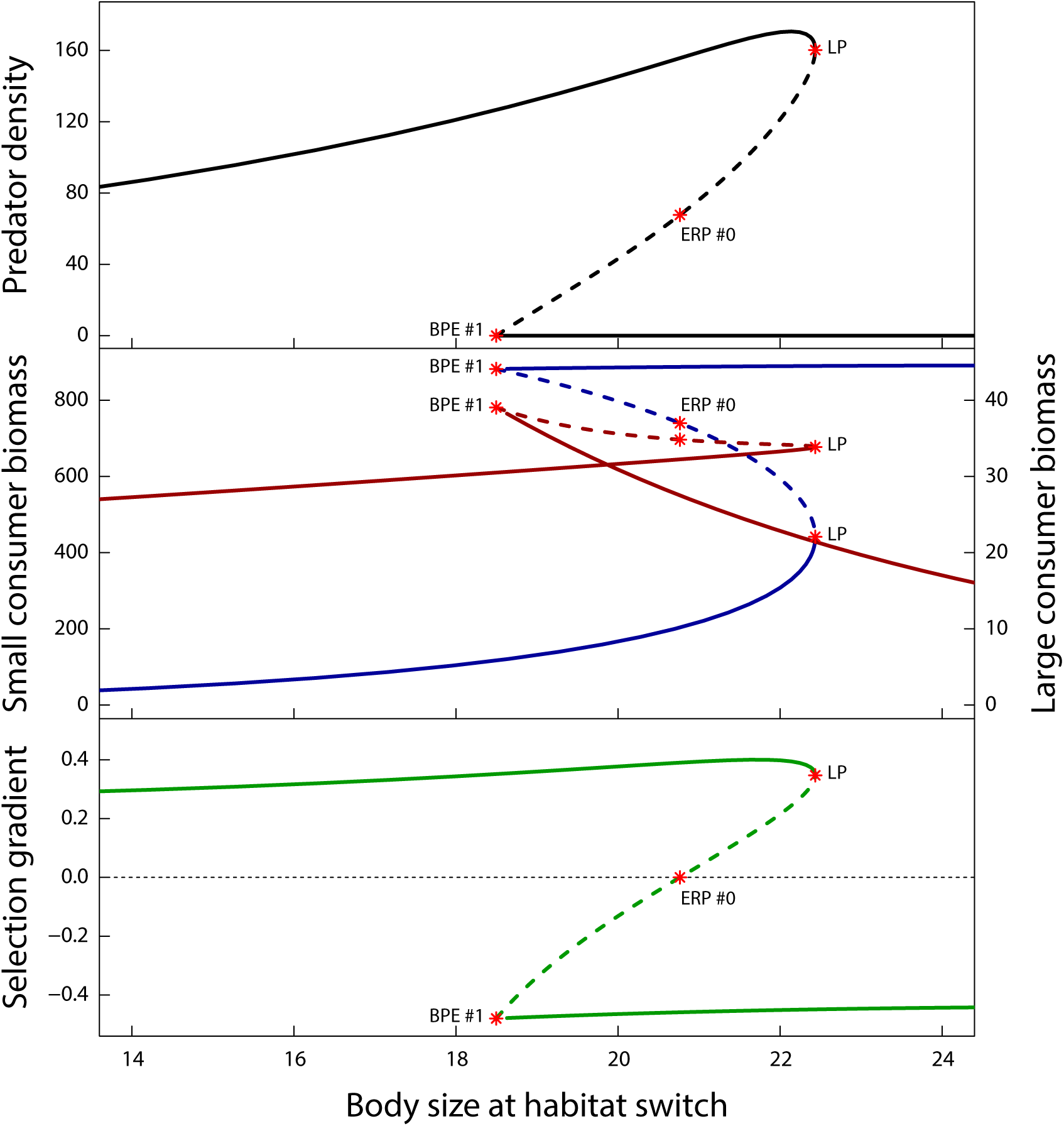
As Figure 3 but for *X*_*max*_ = 5.0. The point labeled `“ERP #0”` refers to an evolutionary repellor for the structured population with index 0 (see main text for details). Notice the opposing signs of the selection gradient in the consumer-resource and predator-consumer-resource equilibrium, which predicts that evolutionary cycling will occur for the body length at habitat switch *ℓ*_*s*_ between the bifurcation points labeled `“BPE #1”` and `“LP”` resulting in repeated invasion and extinction of the predator population.

## Discussion

This paper outlines a general methodology for the analysis of PSPMs, presents a strategy how to apply this methodology to ecological and evolutionary questions and introduces a `R` package that implements the numerical tools required by this methodology. This methodology has been used to gain insight about how individual development affects the ecological dynamics of size-structured populations and communities (de Roos & Persson, 2013). More recently, it was applied to a variety of evolutionary problems, ranging from the evolution of ontogenetic niche shifts (ten Brink & de Roos, 2017), metamorphosis (ten Brink et al., 2019), cannibalism (Hin & de Roos, 2019a), ontogenetic size-scaling (Hin & de Roos, 2019b) and the timing of habitat shifts (Chaparro Pedraza & de Roos, 2020). Given the importance of the individual life history for many eco-evolutionary questions and the importance of environmental feedback on this life history (Lion, 2018), however, there is scope to apply the methodology to a wide range of eco-evolutionary problems.

Physiological structured population models are key to the presented methodology. In contrast to for example integral projection models (Ellner et al., 2016) PSPMs are not directly based on observational life history data. Such data, however, are often collected under at most a few different sets of environmental or density-dependent conditions and hence offer limited information about how the individual life history changes under environmental feedback. In contrast, PSPMs are based on a functional model of the individual life history that mechanistically accounts for the interplay between the life history and the environment (de Roos, 2020). PSPMs therefore often address the question how the individual life history is shaped by both individual traits and environmental, density-dependent impacts. The current paper shows that to address such questions it suffices to specify only the model of the individual life history model dependent on individual traits and environmental variables. Formulating a population-level model is not required as the translation to the population level is sufficiently generic that it can be abstracted into a software approach.

The capabilities of the ‘PSPManalysis’ package are more extensive than highlighted in this paper. Next to the routines for the bifurcation analysis of ecological steady states, the package includes routines for the demographic analysis of PSPMs (following de Roos, 2008), the simulation of the ecological dynamics of structured populations (de Roos et al., 1992), the computation of evolutionary singular states as a function of parameters and the computation of the individual life history under different environmental conditions. All procedures use the same specification of the individual life history, such as the one shown in Table 2, which moreover has a generic structure that is readily adapted to many different life history models. Other methods for the bifurcation analysis of PSPMs do exist (Breda, Diekmann, Gyllenberg, Scarabel, & Vermiglio, 2015; Gyllenberg, Scarabel, & Vermiglio, 2018) and can handle more complex bifurcations, such as Hopf bifurcation points, that can not be detected and analysed with the ‘PSPManalysis’ package, but these methods always have to be specifically tailored to the particular PSPM. The generic and easy-to-adapt implementation of the life history model is the key aspect of the ‘PSPManalysis’ package that makes it useful for a wide range of eco-evolutionary problems.

Theoretical studies in ecology and evolutionary biology often aim for deriving analytical insight, as such insight is considered to apply more generally to a wide range of systems. The complexity of models that can be investigated analytically is, however, severely limited. For example, conditions determining the equilibrium of a size-structured consumer-resource system and its stability can be derived analytically only if mortality is size-independent and an explicit expression for the size-at-age relationship is available (de Roos et al., 1990). The derived conditions are themselves, however, not transparent and nevertheless have to be solved numerically. To facilitate mathematical analysis structured models are then often simplified, for example by replacing size-dependent life history functions with far easier age-dependent ones. Size-dependent and age-dependent life histories (and hence size-structured and age-structured population models) are however fundamentally different from each other, if growth in body size and hence the progression through ontogeny depends on environmental factors like food availability (de Roos & Persson, 2013). Often the choice is hence between a simple model for which analytical results are possible or a model with a more faithful representation of the complexity of real life histories that can only be investigated numerically. In this context the power of numerical bifurcation analysis (Kuznetsov, 1998) has received too little attention. It offers a more powerful approach than brute-force numerical simulations of dynamics, as it provides more comprehensive insight about the different types of stable dynamic patterns that can occur for given combinations of model parameters. The very essence of bifurcation theory furthermore guarantees that these dynamic patterns occur over at least a range of parameter values, lending the results some measure of generality. Methodology and software for the numerical bifurcation analysis of models in terms of ODEs have been available for a while (Dhooge, Govaerts, & Kuznetsov, 2003), the ‘PSPManalysis’ package is intended to provide some of the same capabilities for the general class of PSPMs and thereby facilitate investigating questions about the relationship between complex individual life histories and the dynamics of populations and communities on both ecological and evolutionary time scales.

## Acknowledgements

This work has been supported by funding from the European Research Council under the European Union’s Seventh Framework Programme (FP/2007-2013) / ERC Grant Agreement No. 322814

## Data accessiblity

No experimental or empirical data are used or collected in this study. The ‘PSPManalysis’ package can be installed from CRAN using the command:

~~~
**install**.**packages**(“PSPManalysis”)
~~~

The most recent version can always be installed using the command:

~~~
devtools::**install_**bitbucket(“amderoos**/**PSPManalysis”, subdir = “R**/**”,
                             build**_**vignettes = TRUE)
~~~

The Supplementary Information contains:

1. Table S1 with model parameters and their default values.
2. A discussion of the implementation of the life history model of Chaparro Pedraza and de Roos (2020) in the `R` script called `“Salmon.R”` (also shown in Table 2).
3. A discussion of the `R` commands used to generate the data for the figures in this paper.
4. The `R` script called `“EcoFigures.R”` with the `R` code to generate Figures 1 and 2.
5. The `R` script called `“EvoFigures.R”` with the `R` code to generate Figures 3, 4 and 5.
6. The `C` header file called `“Salmon.h”` with the `C` implementation of the life history model of Chaparro Pedraza and de Roos (2020).

