## Supplementary information for "A general approach for analysis of physiologically structured population models: the R package ‘PSPManalysis’"

### – Supporting Information –

André M. de Roos

Institute for Biodiversity and Ecosystem Dynamics
University of Amsterdam, Amsterdam, The Netherlands

and

Santa Fe Institute, Santa Fe, New Mexico 87501, USA

ORCID ID: https://orcid.org/0000-0002-6944-2048

June 26, 2020

### Contents


**1 Model parameters with their default values** **3**

**2 Implementation in R of the life history model** **4**

**3 R commands to produce the data for all figures** **9**

**4 R script to generate Figure 1 and 2 of the paper** **15**

**5 R script to generate Figure 3, 4 and 5 of the paper** **20**

**6 Implementation in C of the life history model** **29**

### 1 Model parameters with their default values

**Table S1**: Parameters and their default values of the individual life history model of Chaparro Pedraza and de Roos (2020).

| Parameter | Value | Unit | Description |
|---|---|---|---|
| *Life history parameters* | | | |
| $K$ | 1 | $g{\cdot}m^{-3}$ | Half saturation resource density |
| $I_{max}$ | 0.0025 | $g{\cdot}cm^{-2}day^{-1}$ | Maximum ingestion proportionality constant |
| $B_{max}$ | 0.002725 | $cm^{-2}day^{-1}$ | Maximum fecundity proportionality constant |
| $\ell_0$ | 2 | cm | Body size of a newborn |
| $\ell_s$ | 20 | cm | Body size at habitat shift |
| $\ell_m$ | 30 | cm | Body size at maturation |
| $\ell_{inf}$ | 115 | cm | Maximum body size at maximum feeding rate |
| $\xi$ | 0.00051 | $day^{-1}$ | Von Bertalanffy growth rate parameter |
| $\mu_1$ | 0.002 | $day^{-1}$ | Background mortality rate in the habitat 1 |
| $\mu_2$ | 0.006 | $day^{-1}$ | Background mortality rate in the habitat 2 |
| $d$ | 0.75 | – | Exponent in size-dependent predation vulnerability |
| *Parameters related to the environment* | | | |
| $\rho$ | 0.01 | $day^{-1}$ | Resource growth rate |
| $X_{1,max}$ | 5.0 | $g{\cdot}m^{-3}$ | Maximum resource density |
| $\phi$ | 0.001 | $day^{-1}$ | Predator attack rate |
| $\mu_p$ | 0.006 | $day^{-1}$ | Predator mortality rate |

#### 2 Implementation in R of the life history model of Chaparro Pedraza and de Roos (2020) for analysis with the 'PSPManalysis' package.

The following is the content of the file `Salmon.R`, which was used to carry out the computations shown in the figures in the main text of the article and was also shown in Table 2 in the article.

```
PSPMdimensions <- c(PopulationNr = 1, IStateDimension = 2,
                    LifeHistoryStages = 3, ImpactDimension = 5)

EnvironmentState <- c(X = "GENERALODE", P = "PERCAPITARATE")

DefaultParameters <- c(Rho = 0.01, Xmax = 5.0,
                       K = 1.0, Imax = 0.0025, Bmax = 0.002725,
                       L0 = 2.0, Ls = 20.0, Lm = 30.0,  Linf = 115.0,
                       Xi = 0.00051, Mu1 = 0.002, Mu2 = 0.006,
                       D = 0.75, Phi = 0.001, Mup = 0.006)

StateAtBirth <- function(E, pars) {
  with(as.list(c(E, pars)),{
    c(Age = 0.0, Length = L0)
  })
}

LifeStageEndings <- function(lifestage, istate, birthstate, BirthStateNr, E, pars) {
  with(as.list(c(E, pars, istate)),{
    maturation = switch(lifestage, Length - Ls, Length - Lm, -1)
  })
}

LifeHistoryRates <- function(lifestage, istate, birthstate, BirthStateNr, E, pars) {
  with(as.list(c(E, pars, istate)),{
    list(
      development = c(1.0,
                      switch(lifestage, Xi*(Linf*X/(K+X) - Length),
```

```
                              Xi*(Linf - Length), Xi*(Linf - Length))),
        fecundity = switch(lifestage, 0, 0, Bmax*Length^2),
        mortality = switch(lifestage, Mu1, Mu2 + Phi*P*Length^(-D),
                           Mu2 + Phi*P*Length^(-D)),
        impact = switch(lifestage,
                        c(Imax*X/(K+X)*Length^2, 0, Length^3, 0, 0),
                        c(Imax*X/(K+X)*Length^2, Phi*Length^(3-D), 0, Length^3, 0),
                        c(Imax*X/(K+X)*Length^2, Phi*Length^(3-D), 0, 0, Length^3))
      )
  })
}

EnvEqui <- function(I, E, pars) {
  with(as.list(c(E, pars)),{
    c(Rho*(Xmax - X) - I[1], I[2] - Mup)
  })
}
```

The first vector `PSPMdimensions` in the implementation above defines the number of structured populations in the model, the number of state variables with which individuals in the structured populations are characterised, the number of distinct life history stages that individuals pass through during life and the number of impact variables that characterise the impact of an individual on its environment. The example model used in this paper includes only a single structured population and individuals are only characterise by 2 state variables, their length $\ell$ in addition to their age $a$, but the package is sufficiently general to allow for the analysis of models with an arbitrary number of structured populations, in which individuals are characterised by an arbitrary number of state variables. The identification of distinct life history stages in the model is important because at the transitions between these stages the life history dynamics may change discontinuously. For example, in the example model a jump in growth rate occurs when individuals migrate to the growth habitat and a jump in fecundity occurs at maturation. The integration of the ODEs (6) has to stop exactly on reaching such stage transitions to yield accurate results. Distinction of the different life history stages is hence necessary for computational reasons.

The last element of the vector `PSPMdimensions` specifies the dimensionality of the impact of an individual on its environment. In the example model an individual impacts its environment in two different ways: by foraging on the resource and by providing food for the predator, corresponding to the functions $I_1(a,\tilde{X},\tilde{P})$ and $I_2(a,\tilde{X},\tilde{P})$ in the systems of life history ODEs (6). The dimensionality of the individual impact on its environment is therefore 2, but in the model implementation this dimensionality is increased to 5 because the values of all impact variables are produced as output by all 'PSPManalysis' computations. Impact variables can hence be added at will to provide output of interest from a model. In particular, in the example model the 3 additional impact variables are to generate as output the total biomass of consumers in the nursery habitat ($\ell < \ell_s$), of immature consumers in the growth habitat ($\ell_s < \ell < \ell_m$) and of adult consumers ($\ell > \ell_m$). These biomass densities were plotted in the middle panels of Figure 1, 3 and 5 in the main text.

The second vector `EnvironmentState` defines the number and names of state variables that characterise the environment of the structured population(s) as well as their type. Variables characterising the environment can be of different type as exemplified by the resource density $X$ and the predator density $P$ in the example model, depending on whether their dynamics is specified by an ODE like for the resource density (type `"GENERALODE"`) or by a per-capita growth rate as is the case for the predator density (type `"PERCAPITARATE"`). The latter designation informs the program that $\tilde{P} = 0$ is a possible equilibrium state for the predator density but $\tilde{R} = 0$ is not a resource steady state. The manual included with the 'PSPManalysis' package discusses also a third type of environment variable (called `"POPULATIONINTEGRAL"`) that is not discussed here.

The last vector of the model implementation is the vector `DefaultParameters` specifying the names and default values of all the parameters occurring in the model. Just like the names of the environment variables, labeling the elements in the default parameter vector with meaningful names facilitates the implementation of the model functions. All routines defining the model equations use `R` constructs like

```
with(as.list(c(E, pars)),{
  ...
})
```

to allow the use of meaningful names of parameters, environmental and individual state variables in the model definition.

The first two functions of the model implementation specify the state at birth of an individual of the structured population and the conditions that determine the transition between different life history stages. The function `StateAtBirth` should return a vector of individual state variables, defining the meaningful names of these state variables as well as their initial value at birth. The function `LifeStageEndings` uses these individual state variables to specify the threshold values that determine the boundaries between life history stages, in the example model $\ell = \ell_s$ and $\ell = \ell_m$. The argument `lifestage` of this function indicates the index of the life history stage that the individual is in at the time of function invocation and can hence be used in a **`switch`**`()` statement to return the value appropriate to specific life history stages. `lifestage` hence takes on a value in the range $1,\ldots,$`LifeHistoryStages`. The function `LifeStageEndings` has two more arguments `birthstate` and `BirthStateNr` that are not used here but play a role in models in which not all individuals are born with the same state at birth. The manual included with the 'PSPManalysis' package discusses such more complicated situations in more detail.

The function `LifeHistoryRates` has to return as a result a list with the elements `development`, `fecundity`, `mortality` and `impact` that specify the value of the life history functions occurring in the right-hand side of the ODEs (6). The list element `development` has to be a vector that specifies the rate of change for all state variables characterising an individual, while the elements `fecundity` and `mortality` are single valued, indicating the reproduction and mortality rate of an individual as a function of its individual state and the environmental variables. Finally, the list element `impact` has to be a vector of length `ImpactDimension` as specified in the vector `PSPMdimensions`. The first and second element of the vector `impact` in the example model represent the foraging rate, $\alpha(\ell,X)$, of individuals in the nursery habitat and the contribution to the predator's food intake by an individual in the growth habitat, equal to $\varepsilon(\ell)$. The third, fourth and fifth element of the vector `impact` are defined equal to $\ell^3$, representing the mass of an individual, but only in case the individual is in the nursery habitat, is in the growth habitat but still immature and when it is mature, respectively. Otherwise these elements are defined equal to 0. These 3 impact functions hence represent the contribution of the individual to the total biomass of consumers in the nursery habitat and to the biomass of immature and mature consumers in the growth habitat, respectively. It should be pointed out that the function `LifeHistoryRates` should not specify the right-hand side of the ODEs (6) but only the life history functions ($\gamma(\ell,X)$, $\beta(\ell)$,

$\mu(\ell,P)$, $\alpha(\ell,X)$ and $\varepsilon(\ell)$ in the example model) occurring in these ODEs. The 'PSPManalysis' package takes care internally of incorporating them into the ODEs to be integrated.

Finally, the function `EnvEqui` has to return a vector with the values of the conditions that determine the steady state of the different environment variables. The order of values in this vector has to correspond to the order of the environment variables. In the example model the two elements of this vector represent the values of the conditions $g\left(\tilde{X}\right) - \tilde{b}I_1(\infty,\tilde{X},\tilde{P})$ and $\tilde{b}I_2(\infty,\tilde{X},\tilde{P}) - \mu_p$, shown in the equilibrium conditions (7). The argument `I` of the function `EnvEqui` represents the population-level impact on the environment. The elements of this vector have already been multiplied by the population birth rate $\tilde{b}$ prior to invoking the function `EnvEqui`. In the example model the arguments `I[1]` and `I[2]` hence correspond to the products $\tilde{b}I_1(\infty,\tilde{X},\tilde{P})$ and $\tilde{b}I_2(\infty,\tilde{X},\tilde{P})$ in the conditions (7).

#### 3 `R` commands to produce the data for all figures

The data for Figure 1 in the main text was computed with the 'PSPManalysis' package using the following commands:

```
# Step 1
EqR   <- PSPMequi(modelname = "Salmon.R", biftype = "EQ",
                  startpoint = c(0.1, 0.1),
                  stepsize =  0.5, parbnds = c(1, 0.01, 10),
                  options = c("popZE", "0", "envZE", "1"))
# Step 2
EqCR  <- PSPMequi(modelname = "Salmon.R", biftype = "EQ",
                  startpoint = c(EqR$bifpoints[1,1:2], 0),
                  stepsize =  0.5, parbnds = c(1, 0.0, 10),
                  options = c("envZE", "1"))
# Step 3
EqPCR <- PSPMequi(modelname = "Salmon.R", biftype = "EQ",
                  startpoint = EqCR$bifpoints[1,1:4],
                  stepsize = -0.5, parbnds = c(1, 0.0, 10))
```

The 3 commands above all use the function `PSPMequi`, the main part of the 'PSPManalysis' package, which takes as first argument the name of the file with the model implementation (the `R` script discussed in the previous section or alternatively the corresponding `C` header file shown in the last section of this supporting information). The function `PSPMequi` first compiles a dynamic library for the problem from the `C` source files included in the package and the model specification file. After successful compilation this dynamic library is invoked to carry out the computations.

The `biftype = "EQ"` argument of `PSPMequi` determines the type of computation to be performed, here indicating that an equilibrium curve should be computed as a function of a single model parameter. The `stepsize` argument indicates whether the curve continuation should start toward higher or lower values of the model parameter and the size of these steps. The first element of the `parbnds` argument indicates which model parameter to vary by specifying the index of this parameter in the vector of parameters, while the second and third element of the `parbnds` argument specify the minimum and maximum value of this parameter at which to stop the computations. Here it is important to point out that, because the 'PSPManalysis' package is written in `C`, specifying indices has to conform to the `C` convention, in which the first element of a vector has index 0, as opposed to `R` in which the first vector element has index 1. The model parameter that is varied during the computation is from here on referred to as the bifurcation parameter.

The 3 commands above differ in their arguments `startpoint` and `options`. The two elements `"popZE"` and `"0"` in the `options` argument of the first command instruct the function to assume a zero equilibrium density for the structured population with index 0 (the structured consumer population in the example model) and the two elements `"envZE"` and `"1"` do the same for the environmental variable with index 1 (the unstructured predator). Indicating that these two populations have zero equilibrium density simplifies computations because it determines a priori the values of some of the unknowns in the equilibrium localisation, while it also allows the program to detect whether or not the computed equilibrium state can be invaded or not by either the structured consumer or the unstructured predator population. Because these two populations have zero density the `startpoint` contains only two values: the value of the bifurcation parameter $X_{max}$ at which to start the computations and an estimate for the only unknown to solve for, $\tilde{X}$. Because of the start in the trivial steady state $(\tilde{b}, \tilde{X}, \tilde{P}) = (0, X_{max}, 0)$ with $X_{max} = 0.1$ the initial value of $\tilde{X}$ in this case can be given exactly.

This first command produces as output a list containing information about the parameter values and numerical settings used to compute the curve (list element `curvedesc`), a matrix containing the data of the computed equilibrium states along the curve (list element `curvepoints`) and two elements that provide information about the detected bifurcation points along the curve (list element `bifpoints`) and their type (list element `biftype`; the 'PSPManalysis' manual gives full details about these output elements). The element `curvepoints` contains next to the value of the bifurcation parameter and the equilibrium values of the unknowns (in the example model $\tilde{b}$, $\tilde{X}$ and $\tilde{P}$) the values of all interaction variables that have been computed for the structured consumer population. It therefore includes the values of the total population biomass of consumers in the nursery and growth habitat, which have been used to construct the middle panel of Figure 1 in the main text. The element `bifpoints` in the output of this command specifies the value of these variables for the single bifurcation point detected along the computed curve. The list element `biftype` labels this points with the string `"BP #0"`, indicating that this bifurcation point represents a branching point (also called transcritical bifurcation point (Kuznetsov, 1998)) for the structured population with index 0 (the consumer population).

Step 2 of the computations uses the bifurcation point located in the first step of the analysis as starting point to compute the curve of consumer-resource steady states as a function of $X_{max}$. A value of 0 for the birth rate $\tilde{b}$ of the structured consumer population is added to the value of $X_{max}$ and $\tilde{X}$ in this bifurcation point (`EqR$bifpoints[1,1:2]`) to complete the starting point of this second step and the pair `"popZE"` and `"0"` is dropped from the option argument as a zero equilibrium density will no longer be enforced for the structured consumer population. The output of this computation is a similar list as was generated by the first computation step. The resulting curve corresponds to the part of the equilibrium curves shown in Figure 1 in the main text with constant resource density, linearly increasing densities of consumer biomass in the nursery and growth habitat and zero density for the unstructured predator. In this curve the `PSPMequi` function detects a branching point (or transcritical bifurcation point) for the environment variable with index 1 (the unstructured predator population), which it labels as `"BPE #1"` (see Figure 1 in the main text). The consumer-resource steady states to the right of this branching point can be invaded by the unstructured predator population, as indicated by the positive per-capita growth rate that the function `PSPMequi` produces as output.

The last step of the analysis uses the detected branching point for the unstructured predator population to start a computation of the steady states with positive predator density as a function of $X_{max}$. The computation of this curve starts off to lower values of $X_{max}$ (notice the negative `stepsize` argument) as otherwise negative predator densities would result. The result of this computation is a folded curve of steady state values which extends to a minimum just below $X_{max} = 4$ and in which the function `PSPMequi` detects a limit point (or saddle-node bifurcation point Kuznetsov, 1998) (labeled `"LP"` in Figure 1 in the main text).

The two curves shown in Figure 2 of the main text have been constructed using the following commands:

```
# Location of BPE #1 as a function of Xmax and Mup
BPEp <- PSPMequi(modelname = "Salmon.R", biftype = "BPE",
                 startpoint = c(EqCR$bifpoints[1,c(1:2,4)], 0.006),
                 stepsize =  0.5, parbnds = c(1, 0.0, 10, 14, 0, 0.05),
                 options = c("envBP", "1"))
```

```
# Location of LP as a function of Xmax and Mup
LPp <-  PSPMequi(modelname = "Salmon.R", biftype = "LP",
                 startpoint = c(EqPCR$bifpoints[1,1:4], 0.006),
                 stepsize =  0.5, parbnds = c(1, 0.0, 10, 14, 0, 0.05))
```

These two commands continue the curves indicating the location of the bifurcation points labeled `"BPE #1"` and `"LP"` shown in Figure 1 in the main text toward higher values of $X_{max}$, while two similar commands have been used to continue these curves to lower values of $X_{max}$ by changing the `stepsize` argument to -0.5. The first command above adds the default predator mortality rate (0.006) to the bifurcation point detected in the consumer-resource equilibrium curve (`EqCR$bifpoints[1,c(1:2,4)]`) to complete the starting point for a computation of type `"BPE"`, which tells the function `PSPMequi` that the type of point to be computed is a branching point of one of the environment variables. The index of the environment variable that the branching point pertains to is indicated with the option pair `"envBP"` and `"1"` in the `options` argument. Similarly, the second command uses the bifurcation point detected in the curve representing the predator-consumer-resource equilibrium (`EqPCR$bifpoints[1,c(1:2,4)]`) for a computation of type `"LP"`, resulting in a curve of limit points. Notice that in both command the `parbnds` arguments now consists of two triplets with the second triplet specifying the index, minimum and maximum value of the second model parameter to vary. Both commands generate a list as output, containing an element `curvepoints` with the data of all computed solution points (see the 'PSPManalysis' manual for details), which has been used to construct Figure 2 in the main text.

The data shown in Figure 3 of the main text have been produced by the following `R` commands:

```
pars <- c(Rho = 0.01, Xmax = 0.5, K = 1.0, Imax = 0.0025, Bmax = 0.002725,
          L0 = 2.0, Ls = 20.0, Lm = 30.0,  Linf = 115.0,
          Xi = 0.00051, Mu1 = 0.002, Mu2 = 0.006,
          D = 0.75, Phi = 0.001, Mup = 0.006)
EvoCR <-  PSPMequi(modelname = "Salmon.R", biftype = "EQ",
                   startpoint = c(25, 3.01798283E-01, 1.05872962E-04),
                   stepsize = -0.1, parbnds = c(6, 5.0, 25), parameters = pars,
                   options = c("envZE", "1", "popEVO", "0"))
EvoPCR <- PSPMequi(modelname = "Salmon.R", biftype = "EQ",
                   startpoint = EvoCR$bifpoints[1,1:4],
                   stepsize = 0.2, parbnds = c(6, 5.0, 25), parameters = pars,
```

```
                    options = c("popEVO", "0"))
```

The first command defines a slightly altered parameter vector with $X_{max} = 0.5$ instead of its default value 5.0. These new parameters take effect by passing them as argument `parameters = pars` to the function `PSPMequi`. The last two commands are similar to the commands to construct Figure 1 in the main text, discussed previously. The starting point of the first computation is taken from the consumer-resource equilibrium curve shown in Figure 1 in the main text. The special aspect of both invocations of `PSPMequi` shown is the option vector `c("popEVO", "0")`, which instructs the program to compute the selection gradient for the bifurcation parameter and that this parameter pertains to the life history of individuals in the structured population with index 0 (i.e. the length-structured consumer population). The element `curvepoints` of the output list generated by these commands now contains an additional column with the value of the selection gradient, that is, the derivative $dR_0(\infty,\tilde{X},\tilde{P})/d\ell_s$ of the lifetime reproductive output $R_0$ with respect to the life-history parameter $\ell_s$ (see bottom panel of Figure 3 in the main text).

The curve separating regions with positive and negative mutant fitness in the PIP of Figure 4 (left panel) was computed using the command:

```
PIPp <- PSPMequi(modelname = "Salmon.R", biftype = "PIP",
                 startpoint = EvoPCR$bifpoints[2,c(1:4,1)],
                 stepsize =  0.1, parbnds = c(6, 2, 15, 6, 2, 15),
                 parameters = pars, options = c("popEVO", "0"))
```

and a similar command with a negative stepsize `stepsize = -0.1`. The `biftype = "PIP"` argument instructs the `PSPMequi` function to compute a boundary with zero fitness dependent on the resident and mutant life-history trait value. The starting point of the computation is the CSS detected in Figure 3 in the main text, to which the value of the life-history parameter $\ell_s$ in the CSS is tagged on for the trait value of the mutant. The computation hence starts in the point where resident and mutant are identical. As for other computations involving two parameters (`"BPE"`, `"LP"`), the argument `parbnds` now contains two triplets, which are however identical as the computation involves the resident and mutant value of the same life-history parameter. For PIP calculations the `options` argument of the function `PSPMequi` has to specify which structured population the life-history parameter applies to.

Finally, the trajectory of $\ell_s$ over evolutionary time shown in Figure 4 (right panel) in the main text was generated using the command:

```
TSevo <- PSPMevodyn(modelname = "Salmon.R",
                    startpoint = c(0.2566152, 35.55643, 0.0006141598, 5.019757),
                    curvepars = c(10,200000), evopars = c(0, 6, 5, 10),
                    parameters = pars)
```

Since the theory of Adaptive Dynamics assumes a separation between the ecological and evolutionary time scales, an evolutionary dynamics simulation always has to start in an ecological equilibrium state. The starting point in the `PSPMevodyn` command shown was taken from the results of the computation shown in Figure 3 in the main text. The `curvepars` argument of `PSPMevodyn` specifies the maximum step size and maximum time on the evolutionary time scale for the integration. The `evopars` argument of `PSPMevodyn` specifies the index of the structured population characterised by the evolving life-history parameter, the index of this parameter and its minimum and maximum value at which to stop the computations (if reached before reaching the maximum integration time).

### 4 R script to generate Figure 1 and 2 of the paper

```
library(PSPManalysis)
if (!exists("par.defaults")) par.defaults <- par(no.readonly = T)

modelfile  <- "Salmon.R"

#####################################################################################
# Equilibrium predator, consumer and resource biomass densities as a function of
# Xmax
EqR   <- PSPMequi(modelname = modelfile, biftype = "EQ",
                  startpoint = c(0.1, 0.1),
                  stepsize =  0.5, parbnds = c(1, 0.01, 10),
                  options = c("popZE", "0", "envZE", "1"))
EqCR  <- PSPMequi(modelname = modelfile, biftype = "EQ",
                  startpoint = c(EqR$bifpoints[1,1:2], 0),
                  stepsize =  0.5, parbnds = c(1, 0.0, 10),
                  options = c("envZE", "1"))
EqPCR <- PSPMequi(modelname = modelfile, biftype = "EQ",
                  startpoint = EqCR$bifpoints[1,1:4],
                  stepsize = -0.5, parbnds = c(1, 0.0, 10))

# Make the plot
stableR   <- (EqR$curvepoints[,1] <= EqR$bifpoints[1,1])
stableCR  <- (EqCR$curvepoints[,1] <= EqCR$bifpoints[1,1])
stablePCR <- (EqPCR$curvepoints[,2] >= EqPCR$bifpoints[1,2])

layout(matrix((1:3), nrow = 3, ncol = 1), heights = c(0.85, 0.85, 1))
par(tcl = 0.5)
par(mar = c(0, 10, 2, 10))
plot(NULL, NULL, type="n", xaxt="n", yaxt="n",
     xlim=c(0.0, 10), ylim = c(0, 200), xlab="", ylab="")
lines(c(0,EqPCR$curvepoints[1,1]), c(0, 0), lwd = 3)
lines(EqPCR$curvepoints[stablePCR,1], EqPCR$curvepoints[stablePCR,3], lwd = 3)
lines(EqPCR$curvepoints[!stablePCR,1], EqPCR$curvepoints[!stablePCR,3],
      lwd = 3, lty = "dashed")
```

```
points(EqCR$bifpoints[,1], EqCR$bifpoints[,3], col="red", pch=8, lwd=2, cex = 2)
text(EqCR$bifpoints[,1], EqCR$bifpoints[,3], EqCR$biftype, pos=4, offset=0.7,
     cex = 1.5)
points(EqPCR$bifpoints[,1], EqPCR$bifpoints[,3], col="red", pch=8, lwd=2,
       cex = 2)
text(EqPCR$bifpoints[,1], EqPCR$bifpoints[,3], EqPCR$biftype, pos=2,
     offset=0.7, cex = 1.5)
axis(1, at = (0:6)*2, labels=F)
axis(2, at = (0:6)*40, labels=T, las=2, cex.axis = 1.6)
mtext("Predator density", 2, line = 6, cex = 1.8)

par(mar = c(0, 10, 0, 10))
plot(NULL, NULL, type="n", xaxt="n", yaxt="n",
     xlim=c(0.0, 10), ylim = c(0, 1300), xlab="", ylab="")
lines(c(0, EqCR$curvepoints[1,1]), c(0, 0), lwd = 3, col = rgb(0, 0, 0.6))
lines(c(0, EqCR$curvepoints[1,1]), c(0, 0), lwd = 3, col = rgb(0.6, 0, 0))
lines(EqCR$curvepoints[stableCR,1], EqCR$curvepoints[stableCR,7], lwd = 3,
      col = rgb(0, 0, 0.6))
lines(EqPCR$curvepoints[stablePCR,1], EqPCR$curvepoints[stablePCR,7],
      lwd = 3, col = rgb(0, 0, 0.6))
lines(EqPCR$curvepoints[!stablePCR,1], EqPCR$curvepoints[!stablePCR,7],
      lwd = 3, col = rgb(0, 0, 0.6), lty = "dashed")
points(EqR$bifpoints[,1], 0, col="red", pch=8, lwd=2, cex = 2)
text(EqR$bifpoints[,1], 0, EqR$biftype, pos=3, offset=0.7, cex = 1.5)
points(EqCR$bifpoints[,1], EqCR$bifpoints[,7], col="red", pch=8, lwd=2, cex = 2)
text(EqCR$bifpoints[,1], EqCR$bifpoints[,7], EqCR$biftype, pos=4, offset=0.7,
     cex = 1.5)
points(EqPCR$bifpoints[,1], EqPCR$bifpoints[,7], col="red", pch=8, lwd=2,
       cex = 2)
text(EqPCR$bifpoints[,1], EqPCR$bifpoints[,7], EqPCR$biftype, pos=1,
     offset=1.0, cex = 1.5)

lines(EqCR$curvepoints[stableCR,1],
      10*(EqCR$curvepoints[stableCR,8]+EqCR$curvepoints[stableCR,9]),
      lwd = 3, col = rgb(0.6, 0, 0))
```

```
lines(EqPCR$curvepoints[stablePCR,1],
      10*(EqPCR$curvepoints[stablePCR,8]+EqPCR$curvepoints[stablePCR,9]),
      lwd = 3, col = rgb(0.6, 0, 0))
lines(EqPCR$curvepoints[!stablePCR,1],
      10*(EqPCR$curvepoints[!stablePCR,8]+EqPCR$curvepoints[!stablePCR,9]),
      lwd = 3, col = rgb(0.6, 0, 0), lty = "dashed")
points(EqCR$bifpoints[,1], 10*(EqCR$bifpoints[,8]+EqCR$bifpoints[,9]),
       col="red", pch=8, lwd=2, cex = 2)
text(EqCR$bifpoints[,1], 10*(EqCR$bifpoints[,8]+EqCR$bifpoints[,9]),
     EqCR$biftype, pos=4, offset=0.7, cex = 1.5)
points(EqPCR$bifpoints[,1], 10*(EqPCR$bifpoints[,8]+EqPCR$bifpoints[,9]),
       col="red", pch=8, lwd=2, cex = 2)
text(EqPCR$bifpoints[,1], 10*(EqPCR$bifpoints[,8]+EqPCR$bifpoints[,9]),
     EqPCR$biftype, pos=3, offset=1.0, cex = 1.5)
axis(1, at = (0:6)*2, labels=F)
axis(2, at = (0:6)*200, labels=T, las=2, cex.axis = 1.6)
axis(4, at = (0:6)*200, labels=(0:6)*10, las=2, cex.axis = 1.6)
mtext("Small consumer biomass", 2, line = 6, cex = 1.8)
mtext("Large consumer biomass", 4, line = 6, cex = 1.8)

par(mar = c(10, 10, 0, 10))
plot(NULL, NULL, type="n", xaxt="n", yaxt="n",
     xlim=c(0.0, 10), ylim = c(0, 9), xlab="", ylab="")
lines(c(0, EqR$curvepoints[stableR,1]), c(0,EqR$curvepoints[stableR,2]),
      lwd = 3, col = rgb(0, 0.6, 0))
lines(EqCR$curvepoints[stableCR,1], EqCR$curvepoints[stableCR,2],
      lwd = 3, col = rgb(0, 0.6, 0))
lines(EqPCR$curvepoints[stablePCR,1], EqPCR$curvepoints[stablePCR,2],
      lwd = 3, col = rgb(0, 0.6, 0))
lines(EqPCR$curvepoints[!stablePCR,1], EqPCR$curvepoints[!stablePCR,2],
      lwd = 3, col = rgb(0, 0.6, 0), lty = "dashed")
points(EqR$bifpoints[,1], EqR$bifpoints[,2], col="red", pch=8, lwd=2, cex = 2)
text(EqR$bifpoints[,1], EqR$bifpoints[,2], EqR$biftype, pos=3, offset=0.7,
     cex = 1.5)
points(EqCR$bifpoints[,1], EqCR$bifpoints[,2], col="red", pch=8, lwd=2,
```

```
        cex = 2)
text(EqCR$bifpoints[,1], EqCR$bifpoints[,2], EqCR$biftype, pos=4, offset=0.7,
     cex = 1.5)
points(EqPCR$bifpoints[,1], EqPCR$bifpoints[,2], col="red", pch=8, lwd=2, cex = 2)
text(EqPCR$bifpoints[,1], EqPCR$bifpoints[,2], EqPCR$biftype, pos=3,
     offset=1.0, cex = 1.5)
axis(1, at = (0:6)*2, labels=T, cex.axis = 1.6)
axis(2, at = (0:6)*2, labels=T, las=2, cex.axis = 1.6)
mtext("Maximum resource density", 1, line = 6, cex = 1.8)
mtext("Resource biomass", 2, line = 6, cex = 1.8)

##################################################################################
# Two parameter plot of LP and BPE as a function of Xmax and Mup

BPEm <- PSPMequi(modelname = modelfile, biftype = "BPE",
                 startpoint = c(EqCR$bifpoints[1,c(1:2,4)], 0.006),
                 stepsize = -0.5, parbnds = c(1, 0.0, 10, 14, 0, 0.05),
                 options = c("envBP", "1"))
BPEp <- PSPMequi(modelname = modelfile, biftype = "BPE",
                 startpoint = c(EqCR$bifpoints[1,c(1:2,4)], 0.006),
                 stepsize =  0.5, parbnds = c(1, 0.0, 10, 14, 0, 0.05),
                 options = c("envBP", "1"))
LPm <-  PSPMequi(modelname = modelfile, biftype = "LP",
                 startpoint = c(EqPCR$bifpoints[1,1:4], 0.006),
                 stepsize = -0.5, parbnds = c(1, 0.0, 10, 14, 0, 0.05))
LPp <-  PSPMequi(modelname = modelfile, biftype = "LP",
                 startpoint = c(EqPCR$bifpoints[1,1:4], 0.006),
                 stepsize =  0.5, parbnds = c(1, 0.0, 10, 14, 0, 0.05))

# Make the plot
par(par.defaults)
par(mar = c(6, 10, 1, 10))
plot(NULL, NULL, type = "n", xaxt="n", yaxt="n",
     xlim=c(0.0, 8), ylim = c(0, 0.012), xlab="", ylab="")
lines(BPEm$curvepoints[,1], BPEm$curvepoints[,5], lwd = 3, col = rgb(0, 0, 0.6))
```

```
lines(BPEp$curvepoints[,1], BPEp$curvepoints[,5], lwd = 3, col = rgb(0, 0, 0.6))
lines(LPm$curvepoints[,1], LPm$curvepoints[,5], lwd = 3, col = rgb(0.6, 0, 0))
lines(LPp$curvepoints[,1], LPp$curvepoints[,5], lwd = 3, col = rgb(0.6, 0, 0))
axis(1, at = (0:6)*2, labels=T, cex.axis = 1.6)
axis(2, at = (0:6)*0.002, labels=T, las=2, cex.axis = 1.6)
mtext("Maximum resource density", 1, line = 4, cex = 1.8)
mtext("Predator mortality", 2, line = 6, cex = 1.8)
legend("topleft", c("BPE", "LP"),
       col = c(rgb(0, 0, 0.6), rgb(0.6, 0, 0)), lwd = 3, cex = 1.5)

####################################################################################
# Clean up
PSPMclean("F")
```

### 5 R script to generate Figure 3, 4 and 5 of the paper

```
library(PSPManalysis)
if (!exists("par.defaults")) par.defaults <- par(no.readonly = T)

modelfile  <- "Salmon.R"

###########################################################################################
# Equilibrium predator, consumer and resource biomass densities as a function of
# Ls with Xmax = 0.5
pars <- c(Rho = 0.01, Xmax = 0.5, K = 1.0, Imax = 0.0025, Bmax = 0.002725,
          L0 = 2.0, Ls = 20.0, Lm = 30.0,  Linf = 115.0,
          Xi = 0.00051, Mu1 = 0.002, Mu2 = 0.006,
          D = 0.75, Phi = 0.001, Mup = 0.006)
EvoCR <-  PSPMequi(modelname = "Salmon.R", biftype = "EQ",
                   startpoint = c(25, 3.01798283E-01, 1.05872962E-04),
                   stepsize = -0.1, parbnds = c(6, 5.0, 25), parameters = pars,
                   options = c("envZE", "1", "popEVO", "0"))
EvoPCR <- PSPMequi(modelname = "Salmon.R", biftype = "EQ",
                   startpoint = EvoCR$bifpoints[1,1:4],
                   stepsize = 0.2, parbnds = c(6, 5.0, 25), parameters = pars,
                   options = c("popEVO", "0"))

# Make the plot
stableCR <-  (EvoCR$curvepoints[,1]  >= EvoCR$bifpoints[1,1])
stablePCR <- (EvoPCR$curvepoints[,2] >= EvoPCR$bifpoints[1,2])

layout(matrix((1:3), nrow = 3, ncol = 1), heights = c(0.85, 0.85, 1))
par(tcl = 0.5)
par(mar = c(0, 10, 2, 10))
plot(NULL, NULL, type="n", xaxt="n", yaxt="n",
     xlim=c(5.0, 9), ylim = c(0, 45), xlab="", ylab="")
lines(c(0,EvoPCR$curvepoints[1,1]), c(0, 0), lwd = 3, lty = "dashed")
lines(c(EvoPCR$curvepoints[1,1], 25), c(0, 0), lwd = 3)
lines(EvoPCR$curvepoints[stablePCR,1], EvoPCR$curvepoints[stablePCR,3], lwd = 3)
lines(EvoPCR$curvepoints[!stablePCR,1], EvoPCR$curvepoints[!stablePCR,3],
```

```
      lwd = 3, lty = "dashed")
points(EvoCR$bifpoints[,1], EvoCR$bifpoints[,3], col="red", pch=8, lwd=2, cex = 2)
text(EvoCR$bifpoints[,1], EvoCR$bifpoints[,3], EvoCR$biftype,
     pos=3, offset=1.5, cex = 1.5)
points(EvoPCR$bifpoints[1,1], EvoPCR$bifpoints[1,3], col="red", pch=8,
       lwd=2, cex = 2)
text(EvoPCR$bifpoints[1,1], EvoPCR$bifpoints[1,3], EvoPCR$biftype[1],
     pos=2, offset=0.9, cex = 1.5)
points(EvoPCR$bifpoints[2,1], EvoPCR$bifpoints[2,3], col="red", pch=8,
       lwd=2, cex = 2)
text(EvoPCR$bifpoints[2,1], EvoPCR$bifpoints[2,3], EvoPCR$biftype[2],
     pos=3, offset=0.9, cex = 1.5)
axis(1, at = (0:10)*1, labels=F)
axis(2, at = (0:6)*10, labels=T, las=2, cex.axis = 1.6)
mtext("Predator density", 2, line = 6, cex = 1.8)

par(mar = c(0, 10, 0, 10))
plot(NULL, NULL, type="n", xaxt="n", yaxt="n",
     xlim=c(5.0, 9), ylim = c(0, 75), xlab="", ylab="")
lines(EvoCR$curvepoints[stableCR,1], EvoCR$curvepoints[stableCR,7],
      lwd = 3, col = rgb(0, 0, 0.6))
lines(EvoPCR$curvepoints[stablePCR,1], EvoPCR$curvepoints[stablePCR,7],
      lwd = 3, col = rgb(0, 0, 0.6))
lines(EvoPCR$curvepoints[!stablePCR,1], EvoPCR$curvepoints[!stablePCR,7],
      lwd = 3, col = rgb(0, 0, 0.6), lty = "dashed")
points(EvoCR$bifpoints[,1], EvoCR$bifpoints[,7], col="red", pch=8, lwd=2, cex = 2)
text(EvoCR$bifpoints[,1], EvoCR$bifpoints[,7], EvoCR$biftype, pos=2,
     offset=1.0, cex = 1.5)

points(EvoPCR$bifpoints[1,1], EvoPCR$bifpoints[1,7], col="red", pch=8,
       lwd=2, cex = 2)
text(EvoPCR$bifpoints[1,1], EvoPCR$bifpoints[1,7], EvoPCR$biftype[1],
     pos=4, offset=1.3, cex = 1.5)
points(EvoPCR$bifpoints[2,1], EvoPCR$bifpoints[2,7], col="red", pch=8,
       lwd=2, cex = 2)
```

```
text(EvoPCR$bifpoints[2,1], EvoPCR$bifpoints[2,7], EvoPCR$biftype[2],
     pos=1, offset=1.3, cex = 1.5)

lines(EvoCR$curvepoints[stableCR,1],
      0.7*(EvoCR$curvepoints[stableCR,8]+EvoCR$curvepoints[stableCR,9]),
      lwd = 3, col = rgb(0.6, 0, 0))
lines(EvoPCR$curvepoints[stablePCR,1],
      0.7*(EvoPCR$curvepoints[stablePCR,8]+EvoPCR$curvepoints[stablePCR,9]),
      lwd = 3, col = rgb(0.6, 0, 0))
lines(EvoPCR$curvepoints[!stablePCR,1],
      0.7*(EvoPCR$curvepoints[!stablePCR,8]+EvoPCR$curvepoints[!stablePCR,9]),
      lwd = 3, col = rgb(0.6, 0, 0), lty = "dashed")
points(EvoPCR$bifpoints[,1], 0.7*(EvoPCR$bifpoints[,8]+EvoPCR$bifpoints[,9]),
       col="red", pch=8, lwd=2, cex = 2)
text(EvoPCR$bifpoints[1,1], 0.7*(EvoPCR$bifpoints[1,8]+EvoPCR$bifpoints[1,9]),
     EvoPCR$biftype[1], pos=1, offset=1.0, cex = 1.5)
text(EvoPCR$bifpoints[2,1], 0.7*(EvoPCR$bifpoints[2,8]+EvoPCR$bifpoints[2,9]),
     EvoPCR$biftype[2], pos=3, offset=1.0, cex = 1.5)
points(EvoCR$bifpoints[,1], 0.7*(EvoCR$bifpoints[,8]+EvoCR$bifpoints[,9]),
       col="red", pch=8, lwd=2, cex = 2)
text(EvoCR$bifpoints[,1], 0.7*(EvoCR$bifpoints[,8]+EvoCR$bifpoints[,9]),
     EvoCR$biftype, pos=2, offset=1.0, cex = 1.5)
axis(1, at = (0:10)*1, labels=F)
axis(2, at = (0:6)*20, labels=T, las=2, cex.axis = 1.6)
axis(4, at = (0:6)*20*0.7, labels=(0:6)*20, las=2, cex.axis = 1.6)
mtext("Small consumer biomass", 2, line = 6, cex = 1.8)
mtext("Large consumer biomass", 4, line = 6, cex = 1.8)

par(mar = c(10, 10, 0, 10))
plot(NULL, NULL, type="n", xaxt="n", yaxt="n",
     xlim=c(5.0, 9), ylim = c(-1.2, 0.3), xlab="", ylab="")
lines(EvoCR$curvepoints[stableCR,1], EvoCR$curvepoints[stableCR,12],
      lwd = 3, col = rgb(0, 0.6, 0))
lines(EvoPCR$curvepoints[stablePCR,1], EvoPCR$curvepoints[stablePCR,12],
      lwd = 3, col = rgb(0, 0.6, 0))
```

```
lines(EvoPCR$curvepoints[!stablePCR,1], EvoPCR$curvepoints[!stablePCR,12],
      lwd = 3, col = rgb(0, 0.6, 0), lty = "dashed")
lines(par("usr")[1:2], c(0,0), lwd = 1, lty = "dashed")
points(EvoCR$bifpoints[,1], EvoCR$bifpoints[,12], col="red", pch=8,
       lwd=2, cex = 2)
text(EvoCR$bifpoints[,1], EvoCR$bifpoints[,12], EvoCR$biftype, pos=2,
     offset=1.0, cex = 1.5)
points(EvoPCR$bifpoints[,1], EvoPCR$bifpoints[,12], col="red", pch=8,
       lwd=2, cex = 2)
text(EvoPCR$bifpoints[1,1], EvoPCR$bifpoints[1,12], EvoPCR$biftype[1],
     pos=2, offset=1.0, cex = 1.5)
text(EvoPCR$bifpoints[2,1], EvoPCR$bifpoints[2,12], EvoPCR$biftype[2],
     pos=1, offset=1.0, cex = 1.5)
axis(1, at = (0:10)*1, labels=T, cex.axis = 1.6)
axis(2, at = -1.2 + (0:3)*0.4, labels=c("-1.2", "-0.8", "-0.4", "0"),
     las=2, cex.axis = 1.6)
mtext("Body size at habitat switch", 1, line = 6, cex = 1.8)
mtext("Selection gradient\n(dR0/dLs)", 2, line = 5, cex = 1.8)


##################################################################################
# PIP as a function of resident and mutant Ls value for Xmax = 0.5

PIPp <- PSPMequi(modelname = modelfile, biftype = "PIP",
                 startpoint = EvoPCR$bifpoints[2,c(1:4,1)],
                 stepsize =  0.1, parbnds = c(6, 2, 15, 6, 2, 15),
                 parameters = pars, options = c("popEVO", "0"))
PIPm <- PSPMequi(modelname = modelfile, biftype = "PIP",
                 startpoint = EvoPCR$bifpoints[2,c(1:4,1)],
                 stepsize = -0.1, parbnds = c(6, 2, 15, 6, 2, 15),
                 parameters = pars, options = c("popEVO", "0"))

# Make the plot
par(par.defaults)
par(mar = c(6, 10, 1, 10))
```

```
plot(NULL, NULL, type = "n", xaxt="n", yaxt="n",
     xlim=c(4, 9), ylim = c(4, 9), xlab="", ylab="")
lines(PIPm$curvepoints[,1], PIPm$curvepoints[,5], lwd = 3, col = rgb(0, 0, 0.6))
lines(PIPp$curvepoints[,1], PIPp$curvepoints[,5], lwd = 3, col = rgb(0, 0, 0.6))
lines(par("usr")[1:2], par("usr")[3:4], lwd = 1, col = "black", lty = "solid")
axis(1, at = (0:10)*1, labels=T, cex.axis = 1.6)
axis(2, at = (0:10)*1, labels=T, las=2, cex.axis = 1.6)
mtext("Resident Ls value", 1, line = 4, cex = 1.8)
mtext("Mutant Ls value", 2, line = 5, cex = 1.8)

##################################################################################
# Evolutionary dynamics of Ls for Xmax = 0.5

TSevo <- PSPMevodyn(modelname = modelfile,
                    startpoint = c(0.2566152, 35.55643, 0.0006141598, 5.019757),
                    curvepars = c(10,200000), evopars = c(0, 6, 5, 10),
                    parameters = pars, options = c("report", "100"))

# Make the plot
layout(matrix((1:3), nrow = 3, ncol = 1), heights = c(0.85, 0.85, 1))
par(tcl = 0.5)
par(mar = c(0, 10, 2, 10))
plot(NULL, NULL, type="n", xaxt="n", yaxt="n",
     xlim=c(0, 8.0E+4), ylim = c(0, 45), xlab="", ylab="")
lines(TSevo$curvepoints[,1], TSevo$curvepoints[,3], lwd = 3)
axis(1, at = (0:10)*1.0E+4, labels=F)
axis(2, at = (0:6)*10, labels=T, las=2, cex.axis = 1.6)
mtext("Predator density", 2, line = 6, cex = 1.8)

par(mar = c(0, 10, 0, 10))
plot(NULL, NULL, type="n", xaxt="n", yaxt="n",
     xlim=c(0, 8.0E+4), ylim = c(0, 65), xlab="", ylab="")
lines(TSevo$curvepoints[,1], TSevo$curvepoints[,7], lwd = 3, col = rgb(0, 0, 0.6))
lines(TSevo$curvepoints[,1], (TSevo$curvepoints[,8]+TSevo$curvepoints[,9]),
      lwd = 3, col = rgb(0.6, 0, 0))
```

```
axis(1, at = (0:10)*1.0E+4, labels=F)
axis(2, at = (0:6)*20, labels=T, las=2, cex.axis = 1.6)
axis(4, at = (0:6)*20, labels=(0:6)*20, las=2, cex.axis = 1.6)
mtext("Small consumer biomass", 2, line = 6, cex = 1.8)
mtext("Large consumer biomass", 4, line = 6, cex = 1.8)

par(mar = c(10, 10, 0, 10))
plot(NULL, NULL, type="n", xaxt="n", yaxt="n",
     xlim=c(0, 8.0E+4), ylim = c(4.8, 6.3), xlab="", ylab="")
lines(TSevo$curvepoints[,1], TSevo$curvepoints[,5], lwd = 3, col = rgb(0, 0.6, 0))
lines(par("usr")[1:2], EvoPCR$bifpoints[2,c(1,1)], lwd = 1, lty = "dashed")
axis(1, at = (0:8)*1.0E+4,
     labels=c("0", "10000", "20000", "30000", "40000", "50000",
              "60000", "70000", "80000"), cex.axis = 1.6)
axis(2, at = 5 + (0:10)*0.4, labels=T, las=2, cex.axis = 1.6)
mtext("Evolutionary time", 1, line = 5, cex = 1.8)
mtext("Body size at habitat switch", 2, line = 6, cex = 1.8)

# Plot only the dynamics of Ls over evolutionary time
par(par.defaults)
par(mar = c(6, 10, 1, 10))
plot(NULL, NULL, type="n", xaxt="n", yaxt="n",
     xlim=c(0, 8.0E+4), ylim = c(4.8, 6.3), xlab="", ylab="")
lines(TSevo$curvepoints[,1], TSevo$curvepoints[,5], lwd = 3, col = rgb(0, 0.6, 0))
lines(par("usr")[1:2], EvoPCR$bifpoints[2,c(1,1)], lwd = 1, lty = "dashed")
axis(1, at = (0:4)*2.0E+4,
     labels=c("0", "20000", "40000", "60000", "80000"), cex.axis = 1.6)
axis(2, at = 5 + (0:10)*0.4, labels=T, las=2, cex.axis = 1.6)
mtext("Evolutionary time", 1, line = 4, cex = 1.8)
mtext("Body size at habitat switch", 2, line = 5, cex = 1.8)

#################################################################################
# Equilibrium predator, consumer and resource biomass densities as a function of Ls # with Xmax =

EvoCR2  <- PSPMequi(modelname = modelfile, biftype = "EQ",
```

```r
                    startpoint = c(25, 3.87864289E-01, 1.15174854E-03),
                    stepsize = -0.1, parbnds = c(6, 10.0, 25),
                    options = c("envZE", "1", "popEVO", "0"))
EvoPCR2 <- PSPMequi(modelname = modelfile, biftype = "EQ",
                    startpoint = EvoCR2$bifpoints[1,1:4],
                    stepsize =  0.2, parbnds = c(6, 10.0, 25),
                    options = c("popEVO", "0"))

# Make the plot
stableCR <-  (EvoCR2$curvepoints[,1]  >= EvoCR2$bifpoints[1,1])
stablePCR <- (EvoPCR2$curvepoints[,2] >= EvoPCR2$bifpoints[2,2])

layout(matrix((1:3), nrow = 3, ncol = 1), heights = c(0.85, 0.85, 1))
par(tcl = 0.5)
par(mar = c(0, 10, 2, 10))
plot(NULL, NULL, type="n", xaxt="n", yaxt="n",
     xlim=c(14.0, 24), ylim = c(0, 180), xlab="", ylab="")
lines(c(0,EvoPCR2$curvepoints[1,1]), c(0, 0), lwd = 3, lty = "dashed")
lines(c(EvoPCR2$curvepoints[1,1], 25), c(0, 0), lwd = 3)
lines(EvoPCR2$curvepoints[stablePCR,1], EvoPCR2$curvepoints[stablePCR,3],
      lwd = 3)
lines(EvoPCR2$curvepoints[!stablePCR,1], EvoPCR2$curvepoints[!stablePCR,3],
      lwd = 3, lty = "dashed")
points(EvoCR2$bifpoints[,1], EvoCR2$bifpoints[,3], col="red", pch=8,
       lwd=2, cex = 2)
text(EvoCR2$bifpoints[,1], EvoCR2$bifpoints[,3], EvoCR2$biftype, pos=3,
     offset=1.5, cex = 1.5)
points(EvoPCR2$bifpoints[,1], EvoPCR2$bifpoints[,3], col="red", pch=8,
       lwd=2, cex = 2)
text(EvoPCR2$bifpoints[,1], EvoPCR2$bifpoints[,3], EvoPCR2$biftype, pos=2,
     offset=0.9, cex = 1.5)
axis(1, at = 14 + (0:6)*2, labels=F)
axis(2, at = (0:6)*40, labels=T, las=2, cex.axis = 1.6)
mtext("Predator density", 2, line = 6, cex = 1.8)
```

```
par(mar = c(0, 10, 0, 10))
plot(NULL, NULL, type="n", xaxt="n", yaxt="n",
     xlim=c(14.0, 24), ylim = c(0, 900), xlab="", ylab="")
lines(EvoCR2$curvepoints[stableCR,1], EvoCR2$curvepoints[stableCR,7],
      lwd = 3, col = rgb(0, 0, 0.6))
lines(EvoPCR2$curvepoints[stablePCR,1], EvoPCR2$curvepoints[stablePCR,7],
      lwd = 3, col = rgb(0, 0, 0.6))
lines(EvoPCR2$curvepoints[!stablePCR,1], EvoPCR2$curvepoints[!stablePCR,7],
      lwd = 3, col = rgb(0, 0, 0.6), lty = "dashed")
points(EvoCR2$bifpoints[,1], EvoCR2$bifpoints[,7], col="red", pch=8,
       lwd=2, cex = 2)
text(EvoCR2$bifpoints[,1], EvoCR2$bifpoints[,7], EvoCR2$biftype, pos=2,
     offset=1.0, cex = 1.5)
points(EvoPCR2$bifpoints[,1], EvoPCR2$bifpoints[,7], col="red", pch=8,
       lwd=2, cex = 2)
text(EvoPCR2$bifpoints[,1], EvoPCR2$bifpoints[,7], EvoPCR2$biftype, pos=4,
     offset=1.3, cex = 1.5)

lines(EvoCR2$curvepoints[stableCR,1],
      10*(EvoCR2$curvepoints[stableCR,8]+EvoCR2$curvepoints[stableCR,9]),
      lwd = 3, col = rgb(0.6, 0, 0))
lines(EvoPCR2$curvepoints[stablePCR,1],
      10*(EvoPCR2$curvepoints[stablePCR,8]+EvoPCR2$curvepoints[stablePCR,9]),
      lwd = 3, col = rgb(0.6, 0, 0))
lines(EvoPCR2$curvepoints[!stablePCR,1],
      10*(EvoPCR2$curvepoints[!stablePCR,8]+EvoPCR2$curvepoints[!stablePCR,9]),
      lwd = 3, col = rgb(0.6, 0, 0), lty = "dashed")
points(EvoPCR2$bifpoints[,1], 10*(EvoPCR2$bifpoints[,8]+EvoPCR2$bifpoints[,9]),
       col="red", pch=8, lwd=2, cex = 2)
text(EvoPCR2$bifpoints[1,1], 10*(EvoPCR2$bifpoints[1,8]+EvoPCR2$bifpoints[1,9]),
     EvoPCR2$biftype[1], pos=3, offset=1.0, cex = 1.5)
text(EvoPCR2$bifpoints[2,1], 10*(EvoPCR2$bifpoints[2,8]+EvoPCR2$bifpoints[2,9]),
     EvoPCR2$biftype[2], pos=4, offset=1.0, cex = 1.5)
points(EvoCR2$bifpoints[,1], 10*(EvoCR2$bifpoints[,8]+EvoCR2$bifpoints[,9]),
       col="red", pch=8, lwd=2, cex = 2)
```

```
text(EvoCR2$bifpoints[,1], 10*(EvoCR2$bifpoints[,8]+EvoCR2$bifpoints[,9]),
     EvoCR2$biftype, pos=2, offset=1.0, cex = 1.5)
axis(1, at = 14+ (0:6)*2, labels=F)
axis(2, at = (0:6)*200, labels=T, las=2, cex.axis = 1.6)
axis(4, at = (0:6)*200, labels=(0:6)*10, las=2, cex.axis = 1.6)
mtext("Small consumer biomass", 2, line = 6, cex = 1.8)
mtext("Large consumer biomass", 4, line = 6, cex = 1.8)

par(mar = c(10, 10, 0, 10))
plot(NULL, NULL, type="n", xaxt="n", yaxt="n",
     xlim=c(14.0, 24), ylim = c(-0.5, 0.5), xlab="", ylab="")
lines(EvoCR2$curvepoints[stableCR,1], EvoCR2$curvepoints[stableCR,12],
      lwd = 3, col = rgb(0, 0.6, 0))
lines(EvoPCR2$curvepoints[stablePCR,1], EvoPCR2$curvepoints[stablePCR,12],
      lwd = 3, col = rgb(0, 0.6, 0))
lines(EvoPCR2$curvepoints[!stablePCR,1], EvoPCR2$curvepoints[!stablePCR,12],
      lwd = 3, col = rgb(0, 0.6, 0), lty = "dashed")
lines(par("usr")[1:2], c(0,0), lwd = 1, lty = "dashed")
points(EvoCR2$bifpoints[,1], EvoCR2$bifpoints[,12], col="red", pch=8,
       lwd=2, cex = 2)
text(EvoCR2$bifpoints[,1], EvoCR2$bifpoints[,12], EvoCR2$biftype, pos=2,
     offset=1.0, cex = 1.5)
points(EvoPCR2$bifpoints[,1], EvoPCR2$bifpoints[,12], col="red", pch=8,
       lwd=2, cex = 2)
text(EvoPCR2$bifpoints[,1], EvoPCR2$bifpoints[,12], EvoPCR2$biftype, pos=2,
     offset=1.0, cex = 1.5)
axis(1, at = 14 + (0:6)*2, labels=T, cex.axis = 1.6)
axis(2, at = -0.4 + (0:6)*0.2, labels=T, las=2, cex.axis = 1.6)
mtext("Body size at habitat switch", 1, line = 6, cex = 1.8)
mtext("Selection gradient\n(dR0/dLs)", 2, line = 5, cex = 1.8)

##################################################################################
# Clean up
PSPMclean("F")
```

### 6 Implementation in C of the life history model of Chaparro Pedraza and de Roos (2020) for analysis with the 'PSPManalysis' package.

```
/*
  Salmon.h -  Header file specifying the elementary life-history functions of
              the model analyzed in:

                P. Catalina Chaparro-Pedraza & Andre M. de Roos.
                Density-dependent effects of mortality on the optimal body
                size to shift habitat: Why smaller is better despite increased
                mortality risk.
                Evolution 74-5: 831-841 (2020)

              The model includes a basic resource and a size-structured
              consumer population that switches habitat from a nursery
              habitat to a growth habitat.
              In addition, a size-selective predator forages on consumers
              in the growth habitat. Vulnerability of consumers to predation
              in the growth habitat scales with L^(-D).

  Layout for the i-state variables:

    istate[0][0]  : Age
    istate[0][1]  : Length

  Layout for the environment variables:

    E[0]          : Resource
    E[1]          : Predators

  Layout for the interaction (and output) variables:

    I[0][0]       : Total resource ingestion by the consumer population
    I[0][1]       : Consumer biomass availability for predators
```

```
    I[0][2]       : Total biomass of small juveniles in the nursery habitat
    I[0][3]       : Total biomass of juveniles in the growth habitat
    I[0][4]       : Total adult biomass

  Last modification: AMdR - Jun 17, 2020
*/

/*
 *===========================================================================
 *  SECTION 1: PROBLEM DIMENSIONS, NUMERICAL SETTINGS AND MODEL PARAMETERS
 *===========================================================================
 */
// Dimension settings: Required
#define POPULATION_NR              1
#define STAGES                     3
#define I_STATE_DIM                2
#define ENVIRON_DIM                2
#define INTERACT_DIM               5
#define PARAMETER_NR               15

// Numerical settings: Optional (default values adopted otherwise)
#define MIN_SURVIVAL               1.0E-9      // Minimum individual survival
#define MAX_AGE                    200000      // Absolute maximum individual age

#define DYTOL                      1.0E-7      // Variable tolerance
#define RHSTOL                     1.0E-6      // Function tolerance

#define ALLOWNEGATIVE              0           // Negative solution values allowed?
#define COHORT_NR                  200         // Number of cohorts in state output


// Descriptive names of parameters in parameter array (at least two are required)
char *parameternames[PARAMETER_NR] = {"Rho", "Xmax",   "K", "Imax",  "Bmax",
                                      "L0",  "Ls",     "Lm", "Linf",  "Xi",
                                      "Mu1", "Mu2",    "D", "Alpha", "Mup"};
```

```
// Default values of all parameters
double parameter[PARAMETER_NR] = {0.01, 5.0, 1.0, 0.0025, 0.002725,
                                  2.0, 20.0, 30.0, 115.0, 0.00051,
                                  0.002, 0.006, 0.75, 0.001, 0.006};

// Aliases definitions for all istate variables
#define AGE       istate[0][0]
#define LENGTH    istate[0][1]

// Aliases definitions for all environment variables
#define X         E[0]
#define P         E[1]

// Aliases definitions for all parameters

// Resource in the nursery habitat
#define RHO       parameter[ 0]      // Resource growth rate
#define XMAX      parameter[ 1]      // Maximum resource density

// Population with habitat shift
#define K         parameter[ 2]      // Half saturation resource density
#define IMAX      parameter[ 3]      // Maximum ingestion proportionality constant

#define BMAX      parameter[ 4]      // Maximum fecundity proportionality constant

#define L0        parameter[ 5]      // Body size of a newborn
#define LS        parameter[ 6]      // Body size at the habitat shift
#define LM        parameter[ 7]      // Body size at maturation
#define LINF      parameter[ 8]      // Maximum body size at maximum feeding rate

#define XI        parameter[ 9]      // Von Bertalanffy growth rate parameter

#define MU1       parameter[10]      // Background mortality rate in habitat 1
#define MU2       parameter[11]      // Background mortality rate in habitat 2
```

```
#define D         parameter[12]      // Exponent of size-dependent mortality

#define PHI       parameter[13]      // Scaled predator attack rate
#define MUP       parameter[14]      // Scaled predator mortality rate

/*
 *===========================================================================
 *  SECTION 2: DEFINITION OF THE INDIVIDUAL LIFE HISTORY
 *===========================================================================
 */

/*
 * Specify the number of states at birth for the individuals in all structured
 * populations in the problem in the vector BirthStates[].
 */

void SetBirthStates(int BirthStates[POPULATION_NR], double E[])
{
  BirthStates[0] = 1;

  return;
}


/*
 * Specify all the possible states at birth for all individuals in all
 * structured populations in the problem. BirthStateNr represents the index of
 * the state of birth to be specified. Each state at birth should be a single,
 * constant value for each i-state variable.
 *
 * Notice that the first index of the variable 'istate[][]' refers to the
 * number of the structured population, the second index refers to the
 * number of the individual state variable. The interpretation of the latter
 * is up to the user.
```

```
 */

void StateAtBirth(double *istate[POPULATION_NR], int BirthStateNr, double E[])
{
  AGE    = 0.0;
  LENGTH = L0;

  return;
}



/*
 * Specify the threshold determining the end point of each discrete life
 * stage in individual life history as function of the i-state variables and
 * the individual's state at birth for all populations in every life stage.
 *
 * Notice that the first index of the variable 'istate[][]' refers to the
 * number of the structured population, the second index refers to the
 * number of the individual state variable. The interpretation of the latter
 * is up to the user.
 */

void IntervalLimit(int lifestage[POPULATION_NR], double *istate[POPULATION_NR],
                   double *birthstate[POPULATION_NR], int BirthStateNr, double E[],
                   double limit[POPULATION_NR])
{
  switch (lifestage[0])
    {
      case 0:
        limit[0] = LENGTH - LS;
        break;
      case 1:
        limit[0] = LENGTH - LM;
        break;
    }
```

```
  return;
}


/*
 * Specify the development of individuals as a function of the i-state
 * variables and the individual's state at birth for all populations in every
 * life stage.
 *
 * Notice that the first index of the variables 'istate[][]' and 'development[][]'
 * refers to the number of the structured population, the second index refers
 * to the number of the individual state variable. The interpretation of the
 * latter is up to the user.
 */

void Development(int lifestage[POPULATION_NR], double *istate[POPULATION_NR],
                 double *birthstate[POPULATION_NR], int BirthStateNr, double E[],
                 double development[POPULATION_NR][I_STATE_DIM])
{
  development[0][0] = 1.0;
  if (lifestage[0] == 0)
    development[0][1] = XI * (LINF * X / (K + X) - LENGTH);
  else
    development[0][1] = XI * (LINF - LENGTH);

  return;
}


/*
 * Specify the possible discrete changes (jumps) in the individual state
 * variables when ENTERING the stage specified by 'lifestage[]'.
 *
 * Notice that the first index of the variable 'istate[][]' refers to the
```

```
 * number of the structured population, the second index refers to the
 * number of the individual state variable. The interpretation of the latter
 * is up to the user.
 */

void DiscreteChanges(int lifestage[POPULATION_NR], double *istate[POPULATION_NR],
                     double *birthstate[POPULATION_NR], int BirthStateNr,
                     double E[])
{
  return;
}



/*
 * Specify the fecundity of individuals as a function of the i-state
 * variables and the individual's state at birth for all populations in every
 * life stage.
 *
 * The number of offspring produced has to be specified for every possible
 * state at birth in the variable 'fecundity[][]'. The first index of this
 * variable refers to the number of the structured population, the second
 * index refers to the number of the birth state.
 *
 * Notice that the first index of the variable 'istate[][]' refers to the
 * number of the structured population, the second index refers to the
 * number of the individual state variable. The interpretation of the latter
 * is up to the user.
 */

void Fecundity(int lifestage[POPULATION_NR], double *istate[POPULATION_NR],
               double *birthstate[POPULATION_NR], int BirthStateNr, double E[],
               double *fecundity[POPULATION_NR])
{
  fecundity[0][0] = 0.0;
```

```
  if (lifestage[0] > 1) fecundity[0][0] = BMAX * LENGTH * LENGTH;

  return;
}



/*
 * Specify the mortality of individuals as a function of the i-state
 * variables and the individual's state at birth for all populations in every
 * life stage.
 *
 * Notice that the first index of the variable 'istate[][]' refers to the
 * number of the structured population, the second index refers to the
 * number of the individual state variable. The interpretation of the latter
 * is up to the user.
 */

void Mortality(int lifestage[POPULATION_NR], double *istate[POPULATION_NR],
               double *birthstate[POPULATION_NR], int BirthStateNr, double E[],
               double mortality[POPULATION_NR])
{
  if (lifestage[0] == 0)
    mortality[0] = MU1;
  else
    mortality[0] = MU2 + PHI * P * pow(LENGTH, -D);

  return;
}



/*
 *===========================================================================
 *  SECTION 3: FEEDBACK ON THE ENVIRONMENT
 *===========================================================================
 */
```

```
/*
 * For all the integrals (measures) that occur in interactions of the
 * structured populations with their environments and for all the integrals
 * that should be computed for output purposes (e.g. total juvenile or adult
 * biomass), specify appropriate weighing function dependent on the i-state
 * variables, the individual's state at birth, the environment variables and
 * the current life stage of the individuals. These weighing functions should
 * be specified for all structured populations in the problem. The number of
 * weighing functions is the same for all of them.
 *
 * Notice that the first index of the variables 'istate[][]' and 'impact[][]'
 * refers to the number of the structured population, the second index of the
 * variable 'istate[][]' refers to the number of the individual state variable,
 * while the second index of the variable 'impact[][]' refers to the number of
 * the interaction variable. The interpretation of these second indices is up
 * to the user.
 */

void Impact(int lifestage[POPULATION_NR], double *istate[POPULATION_NR],
            double *birthstate[POPULATION_NR], int BirthStateNr, double E[],
            double impact[POPULATION_NR][INTERACT_DIM])
{
  impact[0][0] = IMAX * X / (K + X) * LENGTH * LENGTH;

  switch (lifestage[0])
    {
      case 0:
        impact[0][1] = 0;
        impact[0][2] = LENGTH*LENGTH*LENGTH;
        impact[0][3] = 0;
        impact[0][4] = 0;
        break;
      case 1:
        impact[0][1] = PHI * pow(LENGTH, -D) * LENGTH * LENGTH * LENGTH;
```

```
        impact[0][2] = 0;
        impact[0][3] = LENGTH*LENGTH*LENGTH;
        impact[0][4] = 0;
        break;
      case 2:
        impact[0][1] = PHI * pow(LENGTH, -D) * LENGTH * LENGTH * LENGTH;
        impact[0][2] = 0;
        impact[0][3] = 0;
        impact[0][4] = LENGTH*LENGTH*LENGTH;
        break;
    }

  return;
}


/*
 * Specify the type of each of the environment variables by setting
 * the entries in EnvironmentType[ENVIRON_DIM] to PERCAPITARATE, GENERALODE
 * or POPULATIONINTEGRAL based on the classification below:
 *
 * Set an entry to PERCAPITARATE if the dynamics of E[j] follow an ODE and 0
 * is a possible equilibrium state of E[j]. The ODE is then of the form
 * dE[j]/dt = P(E,I)*E[j], with P(E,I) the per capita growth rate of E[j].
 * Specify the equilibrium condition as condition[j] = P(E,I), do not include
 * the multiplication with E[j] to allow for detecting and continuing the
 * transcritical bifurcation between the trivial and non-trivial equilibrium.
 *
 * Set an entry to GENERALODE if the dynamics of E[j] follow an ODE and 0 is
 * NOT an equilibrium state of E. The ODE then has a form dE[j]/dt = G(E,I).
 * Specify the equilibrium condition as condition[j] = G(E,I).
 *
 * Set an entry to POPULATIONINTEGRAL if E[j] is a (weighted) integral of the
 * population distribution, representing for example the total population
 * biomass. E[j] then can be expressed as E[j] = I[p][i]. Specify the
```

```
 * equilibrium condition in this case as condition[j] = I[p][i].
 *
 * Notice that the first index of the variable 'I[][]' refers to the
 * number of the structured population, the second index refers to the
 * number of the interaction variable. The interpretation of the latter
 * is up to the user. Also notice that the variable 'condition[j]' should
 * specify the equilibrium condition of environment variable 'E[j]'.
 */

const int EnvironmentType[ENVIRON_DIM] = {GENERALODE, PERCAPITARATE};

void EnvEqui(double E[], double I[POPULATION_NR][INTERACT_DIM],
             double condition[ENVIRON_DIM])
{
  condition[0] = RHO * (XMAX - X) - I[0][0];
  condition[1] = I[0][1] - MUP;

  return;
}


/*==================================================================================*/
```
